# Validated removal of nuclear pseudogenes and sequencing artefacts from mitochondrial metabarcode data

**DOI:** 10.1101/2020.06.17.157347

**Authors:** C. Andújar, T. J. Creedy, P. Arribas, H. López, A. Salces-Castellano, A. Pérez-Delgado, A. P. Vogler, B. C. Emerson

## Abstract

Metabarcoding of Metazoa using mitochondrial genes may be confounded by both the accumulation of PCR and sequencing artefacts and the co-amplification of nuclear mitochondrial pseudogenes (NUMTs). The application of read abundance thresholds and denoising methods is efficient in reducing noise accompanying authentic mitochondrial amplicon sequence variants (ASVs). However, these procedures do not fully account for the complex nature of concomitant sequences and the highly variable DNA contribution of individuals in a metabarcoding sample. We propose, as a complement to denoising, the metabarcoding Multidimensional Abundance Threshold Evaluation (*metaMATE*) framework, a novel approach that allows comprehensive examination of multiple dimensions of abundance filtering and the evaluation of the prevalence of unwanted concomitant sequences in denoised metabarcoding datasets. *metaMATE* requires a denoised set of ASVs as input, and designates a subset of ASVs as being either authentic (mtDNA haplotypes) or non-authentic ASVs (NUMTs and erroneous sequences) by comparison to external reference data and by analysing nucleotide substitution patterns. *metaMATE* (i) facilitates the application of read abundance filtering strategies, which are structured with regard to sequence library and phylogeny and applied for a range of increasing abundance threshold values, and (ii) evaluates their performance by quantifying the prevalence of non-authentic ASVs and the collateral effects on the removal of authentic ASVs. The output from *metaMATE* facilitates decision-making about required filtering stringency and can be used to improve the reliability of intraspecific genetic information derived from metabarcode data. The framework is implemented in the *metaMATE* software, available at https://github.com/tjcreedy/metamate).

## 1 INTRODUCTION

Bulk DNA amplification and high-throughput sequencing (HTS) of biological samples, known as metabarcoding (Taberlet, Coissac, Pompanon, Brochmann, & Willerslev, 2012), is becoming an established tool for the study of biodiversity (e.g., Hamady, Walker, Harris, Gold, & Knight, 2008; Yu et al., 2012). In metabarcoding, authentic amplicon sequence variants (ASVs; Callahan, McMurdie & Holmes 2017) amplified from target genes are inherently accompanied by non-authentic variants. The latter arise from errors accumulated through the amplification and sequencing steps and, in the case of metabarcoding using mitochondrial markers, from the co-amplification of mitochondrial-like templates in the nuclear genome, the so-called nuclear mitochondrial sequences (NUMTS, Lopez *et al.* 1994). Sequence variants derived from heteroplasmy may also be coamplified and sequenced, adding to the complexity of the metabarcoding output. The analysis of community data derived from mitochondrial metabarcoding requires sequences that are free of experimental errors and free of paralogous copies, but how to obtain such ‘clean’ data is a central topic in metabarcoding research. Sequencing reads have frequently been grouped into Operational Taxonomic Units (OTUs) to broadly represent species, but while this process reduces the impact of sequencing noise (R. C. Edgar, 2013; Schloss & Westcott, 2011), it has the undesirable effect of collapsing fine-scale true variants largely precluding the study of intraspecific variation. One option is the application of denoising protocols that have proven efficient in reducing noise accompanying authentic ASVs, such as UNOISE (R. Edgar, 2016), DADA2 (Callahan et al., 2016), or Deblur (Amir et al., 2017), among others. These methods provide the opportunity for direct analysis of sequence variants, without the need for OTU clustering, resulting in improved resolution and reproducibility of metabarcoding data (Callahan et al., 2017).

The efficiency of denoising procedures may be complicated both by variable DNA contributions among individuals in a metabarcoding sample, and the different potential origins of non-authentic ASVs. Patterns of read abundance, similarity to target gene copies, and read-quality scores are generally expected to be more predictable for ASVs derived from PCR and sequencing error, compared to co-amplified NUMTs or heteroplasmic variants. ASVs derived from PCR and sequencing errors typically show low read abundance and only limited divergence from higher frequency authentic ASVs (with the exception of chimeras, for which specific filtering methods have been designed, e.g., Edgar *et al.* 2011; Callahan *et al.* 2016). Additionally, errors generated in the sequencing step are expected to show low base quality scores (Callahan et al., 2016). In most cases, heteroplasmic variants affecting coding regions represent single point mutations, and are present in relatively low abundance compared to the predominant haplotype (Huang et al., 2019; Rensch, Villar, Horvath, Odom, & Flicek, 2016), which can be considered the authentic ASV. NUMTs are also expected to be amplified with lower relative read-abundance compared to authentic ASVs, as the nuclear genome copy number is lower than that of the mitochondrial genome by a factor of 100 to 10000, depending on the taxon, cell type, and tissue (Bogenhagen, 2012; Quiros, Goyal, Jha, & Auwerx, 2017). However, NUMTs, like heteroplasmic variants, are derived from a true genomic template and thus are not expected to show lower quality scores compared to mitochondrial copies. Additionally, the origin and evolutionary dynamics of NUMTs (e.g., Bensasson *et al.* 2001; Hazkani-Covo, Sorek & Graur 2003; Pons & Vogler 2005), means that they can be very divergent from the current mitochondrial genome of an individual. Such divergence is a function of (i) the mutation rate within the nuclear region where NUMT insertion has occurred, (ii) the mutation rate of the mitochondrial genome, and (iii) the time since the nuclear insertion (Bensasson, Zhang, Hartl, & Hewitt, 2001), with some insertions estimated to have occurred as much as 58 million years ago (Bensasson, Feldman, & Petrov, 2003). A subset of NUMTs are easily recognised based on frameshift mutations, in-frame stop codons, or mutational patterns inconsistent with functional genes. Such mutations can quickly accumulate due to the absence of selective forces in the inserted nuclear regions. However, other NUMTs have no obvious features to distinguish them from a mitochondrial copy, as has been documented in DNA barcoding studies (Creedy et al., 2019; Shokralla et al., 2014; Song, Buhay, Whiting, & Crandall, 2008). Among the latter, there may be minor variants of authentic ASVs, representing recent nuclear insertions. However, other NUMTs will retain their functional structure as ancestrally “frozen” pseudogenes (Bensasson et al., 2001), resembling the ancient mitochondrial copy from which they were derived, but with a higher divergence from the current mitochondrial haplotype, by accumulation of changes in the latter.

Given these characteristics of NUMTs, and their well-documented prevalence in most eukaryote taxa (Bensasson et al., 2001; Richly & Leister, 2004), co-amplification of NUMTs may be an important contribution to the high proportion of unexpected sequences found in metabarcoding of mock communities with known haplotype composition (Elbrecht, Vamos, Steinke, & Leese, 2018), in barcoding of single specimens using HTS (e.g., Wang *et al.* 2018; Creedy *et al.* 2019), and also the higher than expected number of OTUs (‘OTU inflation’) found in some metazoan metabarcoding approaches (e.g., Flynn *et al.* 2015; Clare *et al.* 2016; Andújar *et al.* 2018b). In contrast, heteroplasmy is expected to have a limited effect on biodiversity estimates derived from metabarcode data. Heteroplasmic copies typically have low read abundance and high similarity to the predominant haplotype (typically differing by only a single point mutation for coding genes), which together facilitate the filtering, together with sequence noise, by the application of denoising methods. Here we first use several metabarcoding data sets from arthropods to illustrate the problem of NUMT co-amplification. These illustrative examples describe patterns of co-occurrence, relative abundance, and phylogenetic relatedness of ASVs that constitute a mixture of authentic mitochondrial haplotypes, NUMTs, and PCR and sequencing errors, highlighting the difficultiy of denoising such datasets. The application of denoising procedures and read abundance thresholds allow different levels of stringency to be used, which can be adjusted to minimise the survival of non-authentic ASVs. However, in complex metabarcoding mixtures, setting highly stringent filtering thresholds carry the risk that rare but authentic mitochondrial haplotypes may also be removed. Conversely, overly conservative thresholds may result in many non-authentic ASVs not being removed. Critically, there is no method available to evaluate the performance of read filtering and denoising procedures in natural communities of unknown composition with regard to the survival of spurious sequences and/or the removal or authentic target sequences. Thus, filtering stringency is decided blindly.

Here we propose the metabarcoding Multidimensional Abundance Threshold Evaluation (*metaMATE*) framework, a comprehensive examination of multiple dimensions of abundance filtering and the evaluation of the prevalence of unwanted concomitant sequences in denoised metabarcode datasets (Fig. 1 and Supp. Fig. S2). The framework is implemented in the *metaMATE* software, available at https://github.com/tjcreedy/metamate. *metaMATE* requires a denoised set of ASVs as input, and before filtering steps, it designates a subset of ASVs as being either authentic (mtDNA haplotypes) or non-authentic ASVs (NUMTs and sequencing artefacts) by comparison to external reference data and by analysing nucleotide substitution patterns. The software incorporates the application of read abundance filtering strategies, including choices for absolute or relative thresholds, which are applied either to whole datasets, individual libraries, within lineages, within taxa, or combined over multiple of these dimensions. For each filtering strategy, *metaMATE* facilitates the batch application of filtering for a range of abundance threshold values. *metaMATE* evaluates the performance of each filtering exercise by quantifying the prevalence of non-authentic ASVs and the collateral effects on the removal of authentic ASVs. Modifications to filtering criteria may alter these proportions, facilitating the identification of optimal parameters which may vary among data sets due to the characteristics of target taxa and target genes. The output from *metaMATE* also allows different research objectives, for which removing non-authentic and retaining authentic ASVs may have different importance, to guide decision-making about optimal filtering strategy. We demonstrate the utility of the method using metabarcode data from mock and natural communities of various complexity. The results indicate that *metaMATE* should be of general utility for metazoan metabarcoding.

**Figure 1.**
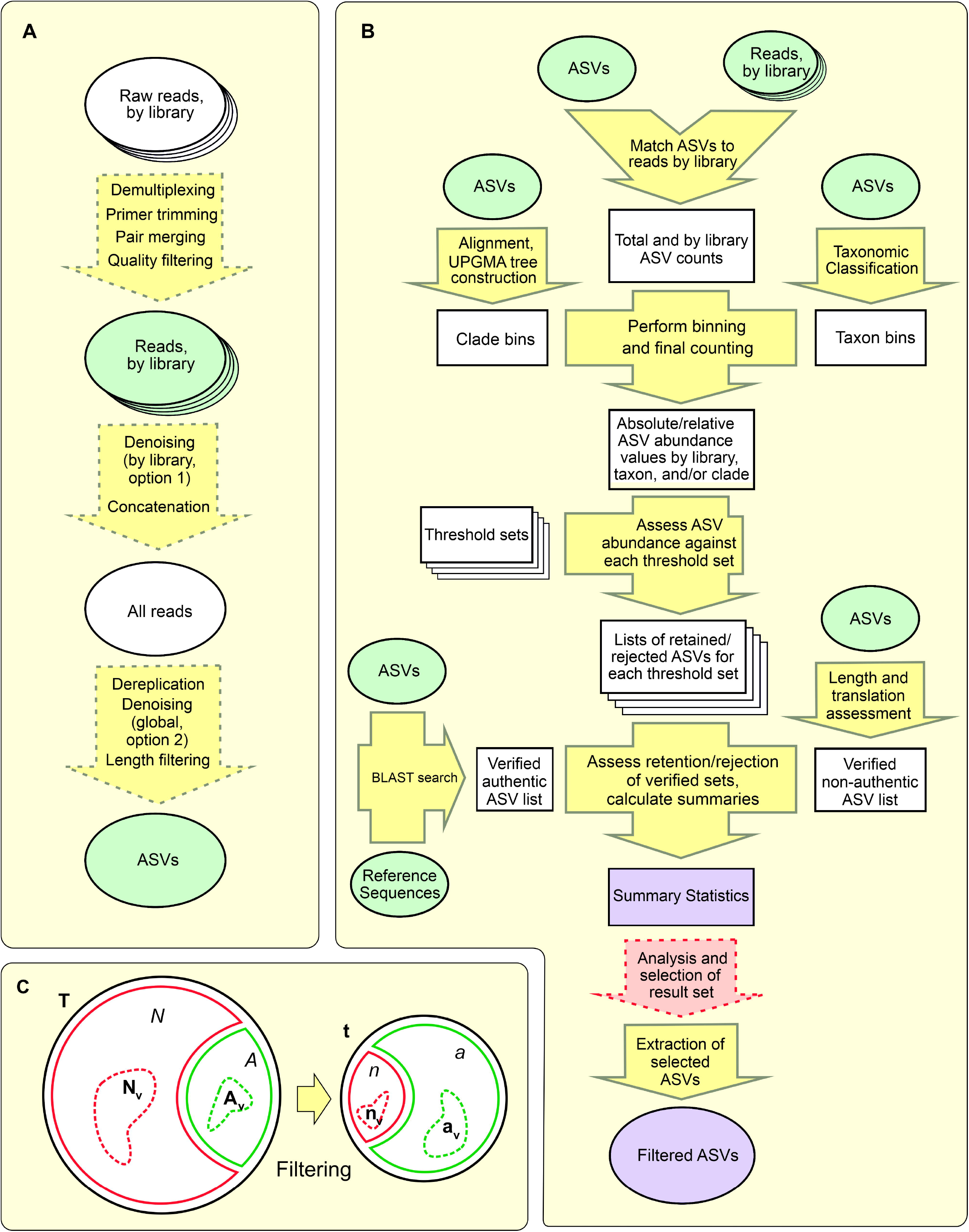
Schematic summaries of generic metabarcode filtering and *metaMATE*. (**A**) Simplified generic metabarcoding pipeline from raw reads to amplified sequence variants (ASVs). (**B**) Key steps and processes within *metaMATE*. Ellipses represent sequence data, rectangles other data, and arrow-shapes processes. Processes conducted externally to *metaMATE* are marked with dashed outline. Stacked ellipses represent multiple parallel data, for example multiple fasta files containing sequences from different libraries. Green indicates input sequences, while purple indicates output from *metaMATE*. Summary statistics describe the performance of different binning strategies and thresholds to simultaneously filter out NUMTs and other erroneous sequences (non-authentic ASVs) while retaining true mitochondrial sequences (authentic ASVs). (**C**). **S**chematic of the expected effect of filtering on the survival of ASVs, where na-ASVs (in red) are removed more effectively than a-ASVs (in gren). *A* and *a* are the numbers of initial and surviving authentic ASVs (a-ASVs); *N* and *n* are the numbers of initial and surviving non-authentic ASVs (na-ASVs); *T* and *t* are the total number of initial and surviving ASVs; the subindex “v” indicates the respective subsets of verified authentic and verified non-authentic ASVs.

## 2 MATERIALS AND METHODS

### 2.1 The *metaMATE* abundance-based ASV filtering and evaluation framework

The *metaMATE* rationale is that read filtering strategies can be evaluated and compared with regard to the prevalence of unwanted concomitant sequences, hereafter referred to as NUMTs (nuclear copies of mitochondrial genes) and noise (variants of target sequences derived from PCR and sequencing error, or heteroplasmic variants with a similar profile, remaining after denoising), versus authentic haplotypes. NUMTs and noise will ideally be removed while retaining the authentic haplotypes. Filtering strategies can be designed to consider read-abundance by dataset, library, or lineage (as a proxy for phylogenetic relatedness), which we implemented in the *metaMATE* program. The same rationale can be used to evaluate other filtering strategies and available denoising methods under a range of stringency parameters. This allows users to decide upon the most appropriate metabarcoding filtering parameters according to their data and research objectives, while providing a measure of the expected number of concomitant sequences in the final dataset. The method comprises two main steps (Fig. 1B):

1. Classification of ASVs. Our starting point is a dataset of ASVs, each of which exclusively belongs to one of two categories: (i) authentic ASVs (a-ASVs) that correspond to actual mitochondrial copies, and (ii) non-authentic ASVs (na-ASVs) that are either noise or NUMTs (see Box 1 for a list of used acronyms). Evaluation of filtering performance relies on the ascertainment of a subset of ASVs that can be confidently considered as authentic sequences (designated as verified-authentic-ASVs; va-ASVs) or non-authentic (designated as verified-non-authentic-ASVs; vna-ASVs). Given the input ASV dataset, va-ASVs can be identified by comparison to reference databases, under the assumption that references are authentic mitochondrial (usually COI barcode) sequences. Any ASV that includes indels and/or stop codons in the translation rendering it non-functional is designated as a vna-ASV. We consider all other ASVs to be unclassified-ASVs (u-ASVs). Pre-processing of the input ASV dataset may vary according to the filtering thresholds to be evaluated, see section 2.2. Improvements in the confident classification of ASVs as va-ASVs or vna-ASVs, such as the availability of local reference databases, will benefit the accuracy of *metaMATE*.
2. Data filtering and evaluation of filtering performance. ASV datasets are subject to filtering procedures with the aim of removing na-ASVs while retaining a-ASVs. Filtering procedures are based on specification of filtering strategies: one or more dimensions by which to evaluate ASV abundance. Each strategy is based on absolute or relative ASV counts either over the entire dataset or by library, optionally further partitioned based on clade and/or taxon membership of ASVs. For each strategy, a range of thresholds can be supplied, and filtering is performed repeatedly for each threshold within each term. For example, the basic strategy employed by most metabarcoding piplines is “minimum absolute total ASV number across the dataset”, usually implemented with a single threshold. The *metaMATE* program can implement this, but can expand to explore the effects of filtering over multiple such thresholds. An alternative strategy would be to filter based on the “minimum absolute number of ASVs per library”, where *metaMATE* would examine the number of reads of an ASV in each library in which it occurred, and reject it if these counts failed to meet the threshold in all libraries in which it occurred. Either of these strategies could alternatively be implemented relative to the total number of reads in the dataset or in each library respectively.

These two basic dimensions of ASV abundance can also be partitioned and implemented in more detailed strategies. The *metaMATE* program can incorporate clade or taxon membership information to filter by, for example, relative ASV abundance by clade within library, whereby the total number of reads for each clade within each library is computed, and then for a given threshold, ASVs are rejected if they fail to meet this threshold for any clade/library combination in which they occur. In total, 16 such combinations of total or library counts, clade and/or taxon membership and absolute or relative counting can be implemented, either individually, successively or synergistically (see below). For each such strategy, a range of thresholds can be specified, and *metaMATE* runs a separate filtering iteration for every threshold for every strategy.

Filtering automatically generates data on the survival of ASVs classified as va-ASV and vna-ASV. The application of a given filtering criterion (e.g. minimum number of reads by library required to retain an ASV) for a range of values for filtering parameters allows trends for the survival of va-ASV and vna-ASV to be analysed.

In addition to obtaining values for the survival of va-ASV and vna-ASV, for each filtering threshold the number of (i) a-ASVs in the initial ASV dataset; (ii) surviving a-ASVs in the filtered dataset; (iii) na-ASVs in the initial dataset, and; (iv) surviving na-ASVs in the filtered dataset can be estimated. These estimations are made from the known values of (i) the number of ASVs in the initial dataset and the number of ASVs retained after filtering and (ii) the proportion of retained va-ASVs and vna-ASVs, under the assumption that va-ASVs and vna-ASVs are a representative subset of all a-ASVs and na-ASVs respectively in the initial dataset. This assumption implies that (i) the ratio between the number of va-ASVs before and after filtering will be similar and can be extrapolated to the ratio between a-ASVs before and after filtering, and (ii) the ratio between the number of vna-ASVs before and after filtering will be similar and can be extrapolated to the ratio between all na-ASVs before and after filtering.

Then, we can estimate the total number of surviving a-ASVs *a* using the formula:

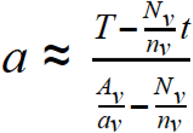

Where *A_v_* and *a*_v_ are the numbers of initial and surviving va-ASVs, *N_v_* and *n_v_* are the numbers of initial and surviving vna-ASVs and *T* and *t* are the total number of initial and surviving ASVs. From this, we can also calculate the estimated total number of surviving na-ASVs *n* and the total number of initial a-ASVs and na-ASVs (A and N) (Fig. 1C; formula derivation in Supplementary Materials).

The *metaMATE* program is written in python3, with some aspects in R. The python3 code utilises the on *biopython* (Cock et al., 2009) library for sequence data handling, and also depends on *numpy* (Oliphant, 2006). The R code makes use of libraries *ape* (Paradis, Claude, & Strimmer, 2004), *phangorn* (Schliep, 2011) and *getopt*. The software allows the user to (i) classify ASVs as va-ASVs and vna-ASVs, (ii) apply any number of filtering criteria constructed from the four dimensions described above alone or in combination, with unlimited customised threshold ranges, facilitating the analysis of trends in survival of va-ASVs and vna-ASVs, and (iii) estimate the number of surviving a-ASV and na-ASVs for each filtering criterion and parameter range explored, using the formula and assumptions described above. Further details on the installation and application of the software are provided in the Supplementary Materials and at https://github.com/tjcreedy/*metamate*.

### 2.2 Empirical application of *metaMATE*

#### 2.2.1 Datasets

Three existing *COI* metabarcoding datasets were used, originating from: (i) 780 individually metabarcoded bees (BEE dataset; Creedy *et al.* 2019) which were used for current purposes to generate 50 *in silico* mock communities of 100 individuals drawn from a subset of 462 confidently identified specimens; (ii) 94 Coleoptera communities from soil samples from the island of Tenerife (COL dataset) (Andújar *et al.* in prep), and (iii) 48 communities of Coleoptera, Acari and Collembola from soil samples from Grazalema, southern Spain (CAC dataset) (Arribas *et al.* in prep) (Suppl. Fig. S1). The three datasets were generated using a nested PCR approach with Nextera XT indexes. In all cases, initial amplification was performed using degenerate primers for 418 bp of the 3’ end of the COI barcode region (Andújar, Arribas, et al., 2018).

To ensure uniform treatment of datasets, raw sequence reads were re-processed following a uniform protocol including primer removal, paired end merging, quality filtering, length filtering for reads ranging between 416-420 bp (the expected 418 bp amplicon ± 2 bp), followed by denoising library by library using UNOISE 3 in USEARCH v11 (Edgar, 2016). The last step included chimera filtering, dereplication, and removal of all singleton reads which were not considered further. The sequences surviving the cleaning and denoising steps (hereafter ASVs) were classified to order or superfamily level, and only ASVs classified as the target taxon or taxa were retained: Apoidea (BEE dataset), Coleoptera (COL) and Coleoptera, Acari and Collembola (CAC). To perform this classification, we generated a reference database comprised of the NCBI *nt* database (downloaded 17 June, 2018) combined with either (i) 1,011 additional reference COI sequences from Coleoptera, Acari and Collembola specimens collected in the Canary Islands and Sierra de Grazalema (COL and CAC datasets) or (ii) the BEEEE reference database described in Creedy et al 2019. For each of the three datasets, searches against the reference database were performed using the BLASTn algorithm (Altschul, Gish, Miller, Myers, & Lipman, 1990), with the following settings: -evalue 0.001, - max_target_seqs 100. Blast results were then processed with MEGAN6 (Huson et al., 2016), using the weighted lowest common ancestor algorithm with default settings to assign taxonomy to ASVs.

#### 2.2.2 Phylogenetic structure, read-abundance, and library co-amplification of ASVs

To illustrate the patterns of library co-amplification, read-abundance, and phylogenetic similarity among noise and NUMTs (na-ASVs) and authentic haplotypes (a-ASVs) generated in *COI* metabarcoding, we conducted maximum-likelihood (ML) phylogenetic analysis to establish relationships of the ASVs within four lineages, and mapped ASV distributions and read abundances against each library using Cytoscape (Shannon et al., 2003). We used two species within the genus *Halictus* and one in the genus *Lasioglossum* (Hymenoptera; Apoidea) from the BEE dataset, and the genus *Cryptocephalus* (Coleoptera: Chrysomelidae) from the COL dataset. A ML tree was first estimated from an alignment (using MAFFT and the FFT-NS-1 algorithm; Katoh & Standley, 2013) of all ASVs from across all libraries for each dataset, with the aim of identifying the clade of ASVs corresponding to the target taxa. All relevant ASVs were extracted, but for logistical purposes related to dataset size and graphical representation, ASVs with a read-count of less than five were excluded from each library and, in the case of *Cryptocephalus,* only the largest 10 libraries (>1000 reads within the *Cryptocephalus* lineage) were selected. ML inferences were conducted in RAxML (Stamatakis, 2006) with 100 searches for the best tree (GTR+G+I model) and 1000 bootstrap pseudoreplicates. In addition, for each dataset a species delimitation analysis was conducted on the ML tree using bPTP (Zhang, Kapli, Pavlidis, & Stamatakis, 2013) on the *bPTP* web server (https://species.h-its.org/) with 100,000 generations and a burn-in of 10%.

#### 2.2.3 Application of *metaMATE*

We applied *metaMATE* to each dataset to identify va-ASVs and vna-ASVs and evaluate the effects on the survival of a-ASVs and na-ASVs under a range of threshold values for different filtering strategies based on read-abundance of ASVs, structured with regard to sequence library and phylogeny. For the COL and CAC datasets, identification of va-ASVs was performed by comparison against the reference set of 1,011 sequences described above combined with the NCBI *nt* database, using BLASTn (-perc_identity 100, only hits with match length >400 bases were considered). For the BEE dataset, identification of va-ASVs was performed by comparison against the set of sequences for the known authentic haplotype of each of the 462 individuals sequenced. All ASVs that included indels and/or stop codons in the translation were designated as vna-ASVs. All others were considered unclassified u-ASVs.

In all cases a given ASV was only excluded if it did not pass the read-abundance threshold in **all**libraries where it was present. The five filtering strategies here tested were: (i) *Absolute ASV abundance by library,* with analysed threshold between 3 and 100 reads. (ii) *Relative ASV abundance by library,* with analysed threshold values between 0.025% to 1%. (iii) *Relative ASV abundance by library **and** 20% similarity clade.* (iv) *Relative ASV abundance by library **and** 15% clade*. (v) *Relative ASV abundance by library **and** 26% clade*. For strategies (iii) to (v), read abundance thresholds from 0.1% to 90% were analysed. Clades were delimited based on specified divergence thresholds from an UPGMA tree of all ASVs constructed using F84 model-corrected distances (Felsenstein & Churchill, 1996) based on a MAFFT FFT-NS-2 alignment (Katoh et al. 2002) of the ASV sequences.

Potential synergy among criteria for removal of na-ASVs while maximizing the survival of a-ASVs was evaluated for the COL dataset, by applying a combination of two of the following strategies: (i) absolute ASV abundance by library, (ii) relative ASV abundance by library, and (iii) relative ASV abundance by library and 20% clade. Only ASVs surviving the application of both criteria were retained.

Finally, using the COL dataset we explored how ASV survival and removal affect: (i) the number of OTUs recovered under similarity thresholds for OTU clustering of 3% and 6%, (ii) the number of surviving OTUs that include one or more va-ASV and consequently can be considered as verified authentic OTUs, and (iii) the number of surviving OTUs that exclusively comprise vna-ASVs and consequently can be considered as verified non-authentic OTUs. The latter contribute to OTU inflation. Similarity clusters were obtained using distances estimated with the F84 model and a UPGMA tree as before.

## 3 RESULTS

Raw sequence reads were subjected to uniform procedures of merging, cleaning, and denoising to establish the total ASVs for each dataset (Fig. 1A), of which a subset could be confidently classified as authentic mitochondrial (va-ASVs) against the respective reference databases (see Material and Methods) or spurious (vna-ASV), leaving all others as unclassified (u-ASVs). The BEE dataset contained 2251 total ASVs identified as Apoidea, including 160 va-ASVs and 117 vna-ASVs. The COL dataset yielded 1845 ASVs classified as Coleoptera, with 74 classified as va-ASVs and 228 as vna-ASVs. The CAC dataset yielded 4804 ASVs, with 712 assigned to Coleoptera (55 va-ASVs and 40 vna-ASVs), 2731 to Acari (99 va-ASVs and 92 vna-ASVs), and 1361 to Collembola (67 va-ASVs and 105 vna-ASVs). Thus, in all cases a large number of AVSs remained “unclassified”.

### 3.1 Phylogenetic structure, read-abundance, and library co-amplification of ASVs

A subset of ASVs was assessed using phylogenetic analysis, to establish the relationships of verified authentic mitochondrial (va-ASVs) and verified non-authentic (vna-ASVs) copies. Phylogenetic relationships and their distributions across libraries were displayed relative to their abundance in a bipartite graph (Fig. 2). The three species of BEE, *Halictus rubicundus* (5 individuals, each sequenced in a separate library), *H. tumulorum* (5 individuals), and *Lasioglossum malachurum* (33 individuals) produced a total of 18, 43, and 45 ASVs, respectively, and included 2, 1 and 2 va-ASVs. In every individual, the most abundant ASV corresponded to a va-ASV (Table 1). A total of and 3, 8 and 8 vna-ASVs were identified for each species respectively, with relatively low read-counts summed across libraries (maximum read count for a vna-ASV = 344; mean for the 19 vna-ASVs = 62) (Table 1). Fifteen of the 19 were shared across 2 or more individuals, including one vna-ASV that was shared across 24 individuals. The 82 unclassified ASVs (u-ASVs) showed relatively low accumulated read-counts summed across libraries (range 10-1795; mean = 87) and 58 of 82 were shared across 2 or more individuals (Fig. 2). Species delimitation analysis on the phylogenetic tree from all ASVs with the bPTP procedure produced two candidate species for both *H. tumulorum* and *L. malachurum*, and in both cases one was exclusively composed of low abundance u-ASVs. It is worth noting that in the case of the BEE dataset all a-ASVs were known, thus all other ASVs had to be either sequencing artefacts or NUMT sequences.

**Figure 2.**
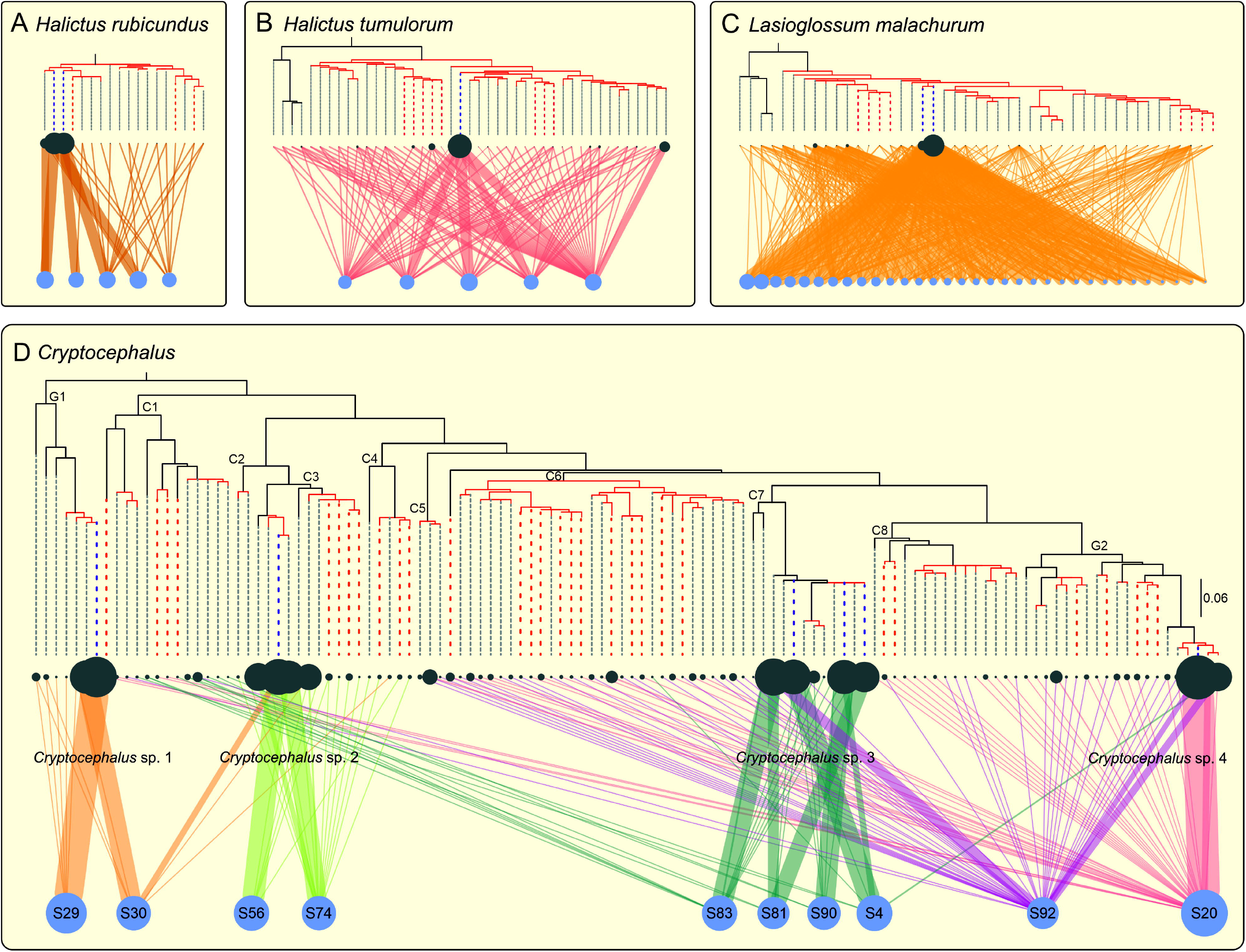
Patterns of phylogenetic relatedness and library co-amplification of ASVs within selected lineages. (**A**) *Halictus rubicundus*, (**B**) *Halictus tumulorum* (**C**), *Lasioglossum malachurum*, and (**D**) *Cryptocephalus*. Graphs show ML phylogenetic trees with mapped distributions of read abundances across libraries (blue circles, with size being proportional to read number) onto each ASV (grey circles, with size being proportional to total abundance across all libraries). Edges of the network represent the presence and abundance (line width) of each ASV within each library. For (A), (B) and (C) each library is a single specimen, whereas in (D) each library includes a complex natural community of beetles where *Cryptocephalus* specimens were present. Clades in red are the best-supported species clusters from bPTP analyses, blue dashed lines highlight ASVs that are identical to a reference sequence (va-ASVs), red dashed lines highlight ASVs with STOP codons or INDELS (vna-ASVs), and grey dashed lines highlight ASVs of unknown origin (u-ASVs). Nodes in (D) are labelled to show clades (C1-C8) and grades (G1-G2) that are exclusively formed by vna-ASVs and u-ASVs.

**Table 1.** Summary of the number of libraries, ASVs, va-ASVs, and vna-ASVS obtained for the three *Halictus* species and the *Crytocephalys* lineage. Read counts refers to the sum of the ASV read-abundance across all libraries where a given ASV is present.

|  | Libraries | ASVs | va-ASVs |  | vna-ASVs |  |
| --- | --- | --- | --- | --- | --- | --- |
|  |  |  | n | read-counts* | n | read counts* |
| **H. rubicundus** | 5 | 18 | 2 | 6915 (6799-7031) | 3 | 15 (10-19) |
| **H. tumulorum** | 5 | 43 | 1 | 8713 (8713-8713) | 8 | 78 (11-344) |
| **L. malachurum** | 33 | 45 | 2 | 48282,5 (4253-92312) | 8 | 64 (11-298) |
| **Cryptocephalus** | 10 | 118 | 6 | 2223 (763-3986) | 30 | 13 (5-43) |

\*mean value (minimum - maximum values)

For the genus *Cryptocephalus,* 6 of 118 ASVs were identified as va-ASVs, each with a high read-count summed across libraries (Fig. 2D, Table 1). Several u-ASVs, closely related to the va-ASVs in the ML tree, showed similarly high read abundances, suggesting their mitochondrial origin (only a subset of the COL authentic haplotypes is known). Several libraries showed more than one high-abundance va-ASV, as expected if more than one *Cryptocephalus* species was present in a sample (libraries correspond to soil samples that sometimes contained tens of *Cryptocephalus* larvae). Thirty ASVs classified as vna-ASVs were found, all of them with low abundances (read-counts summed across libraries from 5 to 43; mean = 13). These vna-ASVs clustered together with additional low abundance u-ASVs into several clades (named C1-C8 in Figure 2D) and grades (named G1 and G2 in Figure 2D). Several of these clades were classified by bPTP as candidate species, producing 39 species in total where only 4 were expected. Despite the notable divergence of sequences inside these clades or grades from the closest va-ASV (e.g, clade C1 has a mean non-corrected p distance of 14.5% against the closest va-ASV), in many cases, vna-ASVs or closely related low abundance u-ASVs were co-amplified in two or more libraries that in addition shared the same *Cryptocephalus* species or closely related species, suggesting a NUMT origin that predates speciation. As an example, libraries S20 and S92 (Fig. 2D) that shared the presence of authentic mitochondrial haplotypes from *Cryptocephalus* sp. 4, also shared one vna-ASV and 3 closely related low abundance u-ASVs within the grade G2 and clade C8. Also, all six libraries including va-ASVs from *Cryptocephalus* sp. 3 and sp. 4 shared co-amplified distantly related ASVs (p-distances >15%) from clade C1 (composed of a set of low abundance u-ASVs and vna-ASVs) (Fig. 2D). In a consistent manner, libraries S29, S30, S56 and S74, that only included va-ASVs from *Cryptocephalus* sp. 1 and sp. 2 did not include any ASVs from clade C1 (Fig. 2D).

### 3.2 Evaluation of filtering efficiency

For all datasets, across all read abundance filtering strategies, increasing thresholds for minimum read abundance resulted in contrasting trends for the removal of va-ASVs and vna-ASVs. In general, the proportion of surviving vna-ASVs dropped quickly below 10% as thresholds were increased, at which point the percentage of surviving va-ASVs exceeded 90% (Fig. 3A, Supp. Fig. S3).

**Figure 3.**
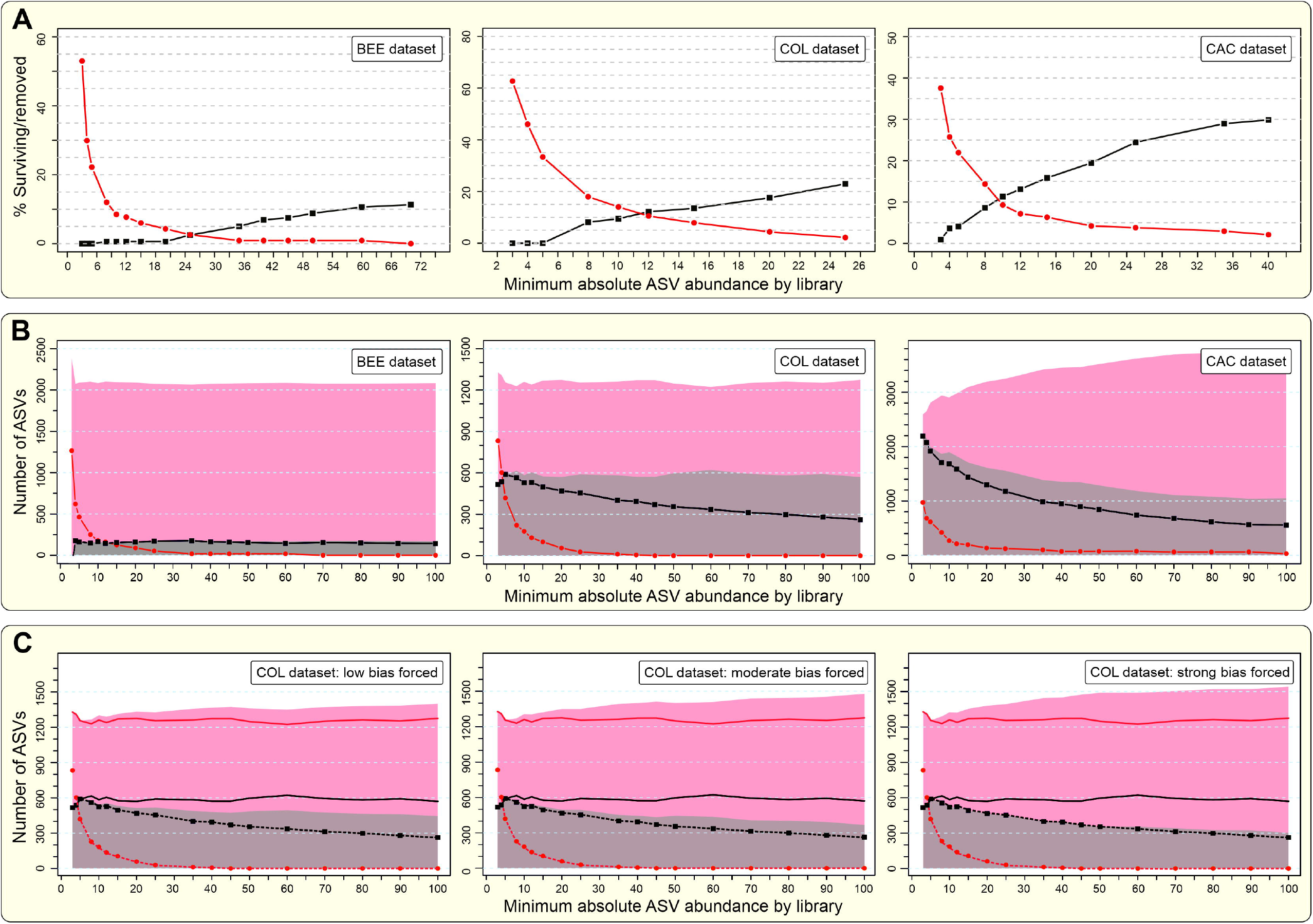
The influence of minimum absolute ASV abundance threshold by library on different measures of filtering success and when sampling is biased against low abundance va-ASVs. (**A**) Proportions of va-ASVs removed (false negatives; black squares and lines) and vna-ASVs retained (false positives; red circles and lines) for three example datasets, bees (BEE), Coleoptera (COL) and Coleptera, Acari and Collembola (CAC). (**B**) Estimated numbers of a-ASVs and na-ASVs comprising initial and filtered ASV datasets for the same three data sets in (A). Grey shading represents the estimated number of initial a-ASVs, red shading represents the estimated number of initial na-ASVs, black squares and lines correspond to estimated number of retained a-ASVs, and red circles and lines correspond to estimated number of retained na-ASVs. (**C**) Estimated number of a-ASVs and na-ASVs in initial and filtered data under low, moderate and strong manipulation to exclude low abundance va-ASVs. Grey and red shading represent the estimated number of initial a-ASVs and initial na-ASVs, respectively. Black squares and red circles represent estimated numbers of retained a-ASVs and retained na-ASVs, respectively. For comparative purposes, red solid, black solid, red dotted, and black dotted lines respectively represent estimations of initial na-ASVs, initial a-ASVs, retained na-ASVs, and retained a-ASVs using the full set of va-ASVs.

The observed values of va-ASVs and vna-ASVs, and estimated values of initial and final a-ASVs and na-ASVs (using the rationale and formula described in methods) are summarized in Figure 3, Supplementary Figures S3 and S4, and Supplementary Tables S1-S15. For the BEE dataset, it was possible to eliminate 99% of vna-ASVs while keeping more than 95% of the va-ASVs using filters for either an absolute or a relative ASV abundance by library. Filtering by the three variants for minimum proportion of reads by similarity cluster and library produced some recalcitrant vna-ASVs that were not removed. For COL and CAC, the filtering criteria generally allowed elimination of 90-95% of vna-ASVs while retaining 80-90% of the va-ASVs. The observed value of vna-ASVs and estimated value of final (surviving) na-ASVs always showed a strong decay reaching 0 (in the case of filtering by minimum absolute or relative ASV abundance by library) or a certain number (filtering based on relative ASV abundance by library and similarity cluster) corresponding to a fixed number of recalcitrant na-ASVs not removable with such criteria. The observed value of va-ASVs and the estimated value of final (surviving) a-ASVs showed a shallower decay with increasing threshold values.

For the BEE and COL datasets, estimates of initial a-ASVs and na-ASVs were approximately constant through increasing threshold values with the strategies of a minimum absolute or relative ASV abundance by library. In the case of BEE, estimated values for *A* and *N* (estimated number of a-ASVs and na-ASVs before filtering) approached the known true values (from Creedy et al. 2019) of *A* = 160 and *N* = 2091. Filtering by minimum absolute ASV abundance by library, we obtained *A* mean = 167 (SD = 10) and *N* mean = 2083 (SD = 10) across different thresholds while minimum relative ASV abundance by library, resulted in *A* mean = 165 (SD = 10) and *N* mean = 2086 (SD = 10). For COL, for the same two criteria, estimated values were always close to *A* = 600 (mean = 581, SD = 20; and mean = 605, SD = 27, respectively) and *N* = 1250 (mean = 1263, SD = 20; and mean = 1238, SD = 27). Estimation of initial a-ASVs and na-ASVs for the CAC dataset showed a different pattern, with a decrease in the estimated value of initial a-ASVs with increasing threshold values, from around 2,200 a-ASVs to 1,000 a-ASVs (mean = 1451, SD = 360; and mean = 1398, SD = 263, respectively for the two criteria), and an increase of initial na-ASVs from 2600 to 3800 (mean = 3245, SD = 360; and mean = 3352, SD = 263). Based on the minimum percentage of reads by similarity clusters, the estimation of final and initial a-ASVs and na-ASVs is less predictable (Supp. Fig. S4), resulting in incorrect values in the case of the mock community (BEE dataset) and stronger trends toward the increasing number of na-ASVs and decreasing number of a-ASVs with increasing threshold values in all datasets.

The simultaneous application of two filtering strategies to the COL dataset improved filtering performance (Table 2). Several of the better filtering combinations resulted in a proportion of surviving vna-ASVs between 2% and 2.6%, estimated to represent 5-6% of all ASVs surviving filtering. The same parameters retained between 82% and 88% of the va-ASVs, estimated as 94-95% of all ASVs surviving filtering. With more stringent combinations of parameters, estimated proportions of na-ASVs in the final dataset can be reduced to 0, 1%, and 2% while still retaining 77%, 80%, and 81% of a-ASVs. (Table 2).

**Table 2.** Filtering performance for a selection of pairwise combinations of filtering criteria and minimum thresholds values for read abundance for the COL dataset. Combinations are shown that minimise the number of surviving verified non-authentic ASVs (vna-ASVs) when the number of excluded verified authentic ASVs (va-ASVs) is between 0 and 17.

| Excl.<br>va-ASVs | Absolute<br>ASV<br>abundance<br>by library | Relative<br>ASV<br>abundance<br>by library | Relative<br>ASV<br>abundance<br>by library<br>and 20%<br>clade | Surviving<br>ASVs<br>(t)** | Surviving<br>va-ASVs<br>(a_v)** | Surviving<br>vna-ASVs<br>(n_v)** | Initial<br>a-ASVs<br>(A)*** | Surviving<br>a-ASV<br>(a)*** | Initial<br>na-ASVs<br>(N)*** | Surviving<br>na-ASV<br>(n)*** | n/t<br>**** |
| --- | --- | --- | --- | --- | --- | --- | --- | --- | --- | --- | --- |
| Pre-filtering initial values |  |  |  | T = 1845 | A<sub>v</sub> = 74 | N<sub>v</sub> = 228 |  |  |  |  |  |
| 0 | 5 | 0.002 | - | 866 (46.9%) | 74 (1000%) | 56 (24.6%) | 547 | 547 | 1298 | 319 | 0.368 |
| 2 | 5 | 0.003 | - | 752 (40.8%) | 72 (97.3%) | 36 (15.8%) | 565 | 550 | 1280 | 202 | 0.269 |
| 3 | 5 |  | 0.015 | 722 (39.1%) | 71 (95.9%) | 28 (12.3%) | 592 | 568 | 1253 | 154 | 0.213 |
| 4 | 5 |  | 0.02 | 685 (37.1%) | 70 (94.6%) | 24 (10.5%) | 584 | 552 | 1261 | 133 | 0.194 |
| 5 | 5 |  | 0.035 | 617 (33.4%) | 69 (93.2%) | 15 (6.6%) | 572 | 533 | 1273 | 84 | 0.136 |
| 6 | - | 0.0035 | 0.035 | 585 (31.7%) | 68 (91.9%) | 14 (6.1%) | 550 | 505 | 1295 | 80 | 0.137 |
| 7 | - | 0.0035 | 0.04 | 572 (31%) | 67 (90.5%) | 14 (6.1%) | 543 | 492 | 1302 | 80 | 0.140 |
| 8 | 5 | - | 0.045 | 535 (29%) | 66 (89.2%) | 11 (4.8%) | 599 | 502 | 1246 | 33 | 0.062 |
| 9 | 8 | - | 0.035 | 556 (30.1%) | 65 (87.8%) | 6 (2.6%) | 596 | 523 | 1249 | 33 | 0.059 |
| 10 | 10 | - | 0.035 | 534 (28.9%) | 64 (86.5%) | 5 (2.2%) | 586 | 506 | 1259 | 28 | 0.052 |
| 11 | 8 | 0.008 | - | 516 (28%) | 63 (85.1%) | 5 (2.2%) | 573 | 488 | 1272 | 28 | 0.054 |
| 12 | 10 | 0.008 | - | 512 (27.8%) | 62 (83.8%) | 5 (2.2%) | 578 | 484 | 1267 | 28 | 0.055 |
| 13 | 10 | - | 0.045 | 513 (27.8%) | 61 (82.4%) | 5 (2.2%) | 589 | 485 | 1256 | 28 | 0.055 |
| 14 | 20 | - | 0.03 | 469 (25.4%) | 60 (81.1%) | 2 (0.9%) | 565 | 458 | 1280 | 11 | 0.023 |
| 15 | 20 | 0.008 | - | 461 (25%) | 59 (79.7%) | 1 (0.4%) | 571 | 455 | 1274 | 6 | 0.013 |
| 17 | 20 | 0.009 | - | 451 (24.4%) | 57 (77.0%) | 0 (0%) | 586 | 451 | 1259 | 0 | 0.000 |

\* Excluded va-ASVs: values lacking (i.e., 1 and 16) represent solutions that were not found with any pairwise combinations of criteria and threshold values; \*\* Observed values; \*\*\* Estimated values; \*\*\*\* The ratio n/t represents the estimated proportion of na-ASVs in the filtered dataset.

Lastly, we examined the effect of increasing filtering stringency on the number of recovered OTUs. Increasing thresholds for minimum read abundance resulted in a similar trend to that found for ASVs, with contrasting results for the removal of those OTUs verified as authentic and non-authentic (Supp. Table S16). The number of surviving OTUs verified as non-authentic (exclusively formed by vna-ASVs) reduces more quickly than the number of OTUs verified as authentic. The proportion of surviving OTUs verified as authentic for both 3% and 6% clustering was very similar to the proportion of surviving va-ASVs for the three filtering criteria and all thresholds values. The proportion of surviving OTUs verified as non-authentic showed a higher rate of survival than that observed for vna-ASVs. As an example, filtering with a minimum relative ASV abundance by library of 0.009 resulted in the survival of 87.8% of va-ASVs, 86.5% of verified authentic OTUs, 4.8% of vna-ASVs, and 21.9% of verified non-authentic OTUs (OTU clustering at 3%) (Supp. Table S16). Thus, “taxonomic inflation” generated by spurious variants can be a more recalcitrant problem than removal of individual NUMTs, requiring higher threshold values for filtering, with an associated cost in the removal of rare species from the dataset.

## 4 DISCUSSION

NUMTs have long been recognised to confound barcoding with Sanger and high throughput sequencing (e.g., Song *et al.* 2008; Shokralla *et al.* 2014; Creedy *et al.* 2019), and the potential impact of NUMTs on metabarcoding has been discussed widely (e.g., Ramirez-Gonzalez *et al.* 2013; Andújar *et al.* 2018b; Elbrecht *et al.* 2018; Liu *et al.* 2019; Delsuc & Ranwez 2020). NUMTs are likely to be consequential in metabarcoding because of: (i) the widespread use of degenerate primers for metabarcoding (e.g. Andújar *et al.* 2018a; Elbrecht *et al.* 2019); (ii) the complexity of specimen mixtures that produce metabarcoding data, and; (iii) the sensitivity of single-molecule sequencing with HTS platforms. NUMT insertions have been documented to occur multiple times within lineages (Bensasson et al., 2003; Hazkani-Covo et al., 2003; Pons & Vogler, 2005; Shi, Dong, Irwin, Zhang, & Mao, 2016), with NUMTs accumulating within genomes over time. In addition, once inserted, duplication events within the nuclear genome may contribute to the formation of NUMT families (Baldo, De Queiroz, Hedin, Hayashi, & Gatesy, 2011; Bensasson et al., 2003; Pamilo, Viljakainen, & Vihavainen, 2007), potentially resulting in hundreds of NUMTs (e.g., Ramos et al., 2011). Critically, NUMTs can retain a fully functional mitochondrial sequence long after their nuclear insertion, if inserted within invariant regions of the nuclear genome (Bensasson et al., 2001). The illustrative examples selected within our metabarcoding datasets highlight the potential magnitude of NUMT diversity in mitochondrial metabarcoding. Despite the difficulty for in fully differentiating NUMTs from noise, ASVs identified as non-authentic (vna-ASVs) exhibiting stop codons and/or frame-shift mutations generally fit patterns of phylogenetic relatedness, read-abundance, co-occurrence, and haplotype sharing across independent libraries that are expected from NUMTs (Fig. 2). The pattern of low-read abundance and library co-amplification of additional ASVs, which are phylogenetically related to vna-ASVs but do not include stop codons and/or frame-shift mutations, point to their probable NUMT origin, illustrating the difficulty to identify all NUMTs exclusively based on their nucleotide sequence.

The accumulation of sequence variants derived from PCR and sequencing error, and the co-amplification of NUMTs, result in a complex mixture of concomitant sequences accompanying target haplotypes in metabarcoding of mitochondrial genes. Read abundance is an obvious filtering parameter. However, read-abundance relationships may be imperfect due to (i) authentic rare haplotypes with relatively low read abundances overlapping with the abundance ranges of concomitant sequences generated from species contributing with more DNA to the DNA pool, and (ii) potential amplification biases increasing the read abundance of some particular NUMT copies. This implies that it is very unlikely that a single abundance threshold can be devised for the removal of all NUMTs and noise sequences from a set of ASVs, while not excluding authentic haplotypes. It also suggests that, because of idiosyncratic heterogeneity and amplification biases, different datasets may vary with regard to optimal criteria and thresholds-parameters to minimise false positives (unwanted sequences retained) and/or false negatives (authentic haplotypes excluded). In this context, denoising procedures are designed to consider not only read abundance, but also similarity among sequences, and even error profiles expected from sequencing technology into their models (e.g. UNOISE, Edgar, 2016; DADA2, Callahan et al., 2016; and Deblur, Amir et al., 2017). However, filtering is typically applied without the possibility of checking performance and deciding upon optimal stringency parameters. Our results show that while denoising likely removes the majority of sequencing errors contributing to ‘noise’, it may be not sufficient to remove all noise and particularly fails to remove sequences that are likely NUMTs. Further filtering is required if the aim is to generate a dataset suitable for haplotype-level analysis, or to reliably eliminate spurious OTUs.

Here we propose a framework to evaluate and select filtering strategies according to user requirements and filtering performance, using a subset of ASVs known to be either authentic mitochondrial haplotypes or undesired NUMTs or noise. We have implemented the evaluation framework within a program that allows for batch application of several filtering strategies under a range of threshold values. We have demonstrated the difficulties for the removal of spurious sequences, and the benefit of evaluating the efficiency of different filtering strategies. *metaMATE* implements several filtering strategies based on absolute read numbers and relative read-abundances of ASVs against the total number of reads in libraries or lineages. For all strategies, opposing trends are observed with increasing threshold value for the removal of verified authentic (va-ASVs) and verified non-authentic (vna-ASVs) sequences. The relative rapid decay of non-authentic sequences allows the elimination of 90-95% of vna-ASVs, while retaining 80-90% of va-ASVs. In addition, *metaMATE* can easily incorporate more complex, custom made filtering strategies. Using paired combinations of filtering criteria, results were improved by removing 98% of vna-ASVs, while retaining 81% of the va-ASVs, and more complex filtering strategies may further improve these results.

After obtaining the survival ratios of both va-ASVs and vna-ASVs, *metaMATE* estimates the number and proportion of surviving a-ASVs and na-ASVs for each abundance threshold (Fig. 1), on the assumption that the subset of va-ASVs and vna-ASVs are representative of the initial number of a-ASVs and na-ASVs, respectively. This allows for the selection of thresholds based on individual acceptance criteria for the maximum number (or proportion) of na-ASVs in a given final dataset. Results from the mock community and real datasets analysed here illustrate the potential utility and issues associated with estimations of a-ASVs and na-ASVs in the initial and final (filtered) datasets. Increasing thresholds based on absolute and relative ASV abundance by library for both the BEE and the COL datasets resulted in estimates of the initial number of a-ASVs (*A*) and na-ASVs (*N)* that are approximately constant, with estimates for BEE approaching the known true values. This supports the reliability of estimates. However, the CAC dataset revealed a different pattern, with a decrease in the estimated number of initial a-ASVs and an increase for initial na-ASVs with increasing threshold values (Supp. Fig. S4). This variation in the estimated values is likely due to the violation of the assumption that va-ASVs are a representative subset of all a-ASVs 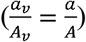, and thus presents a potential means to evaluate the assumption itself. To explore this further, we manipulated the subset of va-ASVs used within the COL dataset to simulate both (i) bias from a lack of low abundance va-ASVs (represented in Fig. 3C), and (ii) bias from a lack of high abundance va-ASVs (Supp. Fig. S5). For both types of bias we explored three intensities: strong, moderate and low. Results reveal that bias generated by a lack of low abundance va-ASVs reproduces the pattern found for the CAC dataset, whereas bias for a lack of high abundance va-ASVs generates the opposite trend. These analyses show that the effect of bias on the estimated initial number of a-ASVs and na-ASVs increases with increasing threshold values. However, they also reveal a limited effect on the estimated number of a-ASVs and na-ASVs in the final dataset, a consequence of the low number of surviving na-ASVs with increasing thresholds. It is also worth noting that filtering strategies based on relative ASV abundance estimated within similarity clusters result in biased estimations, likely due to the prevalence of recalcitrant na-ASVs associated to clusters exclusively formed by a single or several na-ASVs. Taken together, these analyses of bias suggest that: (i) if the assumption of the ratios 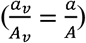 is met, the correct estimation of both initial and final (surviving) numbers of a-ASVs and na-ASVs is straightforward; (ii) violation of the assumption results in predictable changes in the initial number of a-ASVs and na-ASVs, with only limited effect on the estimation of the number of a-ASVs and na-ASVs in the final dataset, and; (iii) estimates obtained from criteria where abundance is calculated within similarity clusters alone are less reliable and should not be used.

Our results also reveal that OTU clustering alone may not be sufficient to remove the effect of na-ASVs. OTUs that are identified as non-authentic can pass filtering based on read-abundance even in higher proportions than individual vna-ASVs. This highlights the problem of “OTU inflation” (Flynn et al. 2015), which we show can be reduced by increasing read-abundance thresholds, but with the expected trade-off for the removal OTUs representing rare species (Supp. Table S16). Thus, the broader *metaMATE* framework can be also used at the OTU level to evaluate filtering performance and the expected taxonomic inflation in datasets before and after filtering, optimising between taxonomic inflation and the removal of rare species.

In conclusion, our results illustrate the presence of NUMT sequences in *COI* metabarcode data and highlight the need to evaluate thresholds for each dataset according to user-defined acceptable levels of false positives and false negatives. Studies seeking data with minimal error, such as for phylogeographic (e.g., Turon et al., 2019) or population genetic analyses (e.g., Elbrecht et al., 2018), should opt for stringent thresholds, to minimise the confounding effect of NUMTs and other spurious sequences, even at the expense of removing some authentic haplotype data and rare species. For other applications, such as those based on measures of beta diversity to explore broad ecological patterns, less strict thresholds may be admissible. In addition, studies aiming to estimate richness values at haplotype or even OTU levels may consider expected biases generated by surviving concomitant sequences to correct data and generate estimates of a-ASVs and authentic OTUs in the initial and final (filtered) datasets. Ultimately these are decisions that can now be made and reported with the incorporation of the proposed evaluation framework in analysis pipelines, by the application of *metaMATE* to ASV datasets generated after denoising. Thus, *metaMATE* builds upon existing denoising strategies to evaluate the reliability of intraspecific genetic information derived from metabarcode data, opening the door for community-level genetic analyses requiring haplotype-level resolution.

## ACKNOWLEDGEMENTS

CA was supported by the Spanish Ministry of Economy and Competitiveness (MINECO, Spain) (CGL2015-74178-JIN) and Fundación CajaCanarias/Obra social “la Caixa”. BCE was supported by the project CGL2017-85718-P (AEI, Spain/FEDER, EU). PA, TJC, BCE and APV were supported by the iBioGen project funded by the H2020 European Research Council, Grant/Award Number: 810729. We extend our gratitude to the regional governments of Andalucía and Canarias (Spain) for facilitating collecting of samples, to Jesús Arribas for assistance with field sampling and Carlos Martínez for the mathematical advice.

## AUTHOR CONTRIBUTIONS

CA and PA conceived and led the study; CA, PA, and TJC designed the methodology; CA, PA, HL, ASC, TPD, APV and BCE provided the data; CA and TJC analysed the data and TJC wrote the *metaMATE* software. CA and BCE wrote the manuscript. All authors contributed critically to the drafts and gave final approval for publication.

## DATA AVAILABILITY

Data will be available from the Dryad Digital Repository upon acceptance

### Box 1. List of acronyms

**ASV**. <u>A</u>mplicon <u>S</u>equence <u>V</u>ariant, sensu Callahan et al. (2016). Each unique DNA sequence within a denoised metabarcoding dataset is an ASV.

**NUMT**. <u>NU</u>clear <u>M</u>i<u>T</u>ocondrial, sensu Lopez et al. 1994. Pseudogenes originating from the insertion of mitochondrial DNA fragments in the nuclear genome.

**a-ASV**. <u>A</u>uthentic <u>A</u>mplicon <u>S</u>equence <u>V</u>ariant. ASV that represents a true sequence amplified from the target gene.

**na-ASV**. <u>N</u>on-<u>a</u>uthentic <u>A</u>mplicon <u>S</u>equence <u>V</u>ariant. ASV with error derived from the PCR and sequencing steps, or by the co-amplification of pseudogenes.

**va-ASV**. <u>V</u>erified <u>a</u>uthentic <u>A</u>mplicon <u>S</u>equence <u>V</u>ariant. ASV that has been verified as authentic by comparison with validated reference sequences.

**vna-ASV**. <u>V</u>erified <u>n</u>on-<u>a</u>uthentic <u>A</u>mplicon <u>S</u>equence <u>V</u>ariant. ASV that has been verified as non-authentic by the presence of mutations incompatible with a functional protein (STOP codons and frame-shift mutations).

**metaMATE**. <u>meta</u>barcoding <u>M</u>ultidimensional <u>A</u>bundance <u>T</u>hreshold <u>E</u>valuation. Approach to evaluate filtering performance, based on the survival of va-ASVs and vna-ASVs, for a range of filtering criteria and thresholds applied to a set of denoised ASVs

## References

Altschul, S. F., Gish, W., Miller, W., Myers, E. W., & Lipman, D. J. (1990). Basic local alignment search tool. Journal of Molecular Biology, 215(3), 403–10. doi:10.1016/S0022-2836(05)80360-2

Amir, A., Daniel, M., Navas-Molina, J., Kopylova, E., Morton, J., Xu, Z. Z., … Knight, R. (2017). Deblur rapidly resolves single-nucleotide community sequence patterns. American Society for Microbiology, 2(2), 1–7. Retrieved from http://genomebiology.biomedcentral.com/articles/10.1186/gb-2012-13-9-r79

Andújar, C., Arribas, P., Gray, C., Bruce, C., Woodward, G., Yu, D. W., & Vogler, A. P. (2018). Metabarcoding of freshwater invertebrates to detect the effects of a pesticide spill. Molecular Ecology, 27(1), 146–166. doi:10.1111/mec.14410

Andújar, C., Emerson, B. C., Arribas, P., Yu, D. W., & Vogler, A. P. (2018). Why the COI barcode should be the community DNA metabarcode for the metazoa, (May), 3968–3975. doi:10.1111/mec.14844

Baldo, L., De Queiroz, A., Hedin, M., Hayashi, C. Y., & Gatesy, J. (2011). Nuclear-mitochondrial sequences as witnesses of past interbreeding and population diversity in the jumping bristletail mesomachilis. Molecular Biology and Evolution, 28(1), 195–210. doi:10.1093/molbev/msq193

Bensasson, D., Feldman, M. W., & Petrov, D. A. (2003). Rates of DNA duplication and mitochondrial DNA insertion in the human genome. Journal of Molecular Evolution, 57(3), 343–354. doi:10.1007/s00239-003-2485-7

Bensasson, D., Zhang, D. X., Hartl, D. L., & Hewitt, G. M. (2001). Mitochondrial pseudogenes: Evolution’s misplaced witnesses. Trends in Ecology and Evolution, 16(6), 314–321. doi:10.1016/S0169-5347(01)02151-6

Bogenhagen, D. F. (2012). Mitochondrial DNA nucleoid structure. Biochimica et Biophysica Acta - Gene Regulatory Mechanisms, 1819(9–10), 914–920. doi:10.1016/j.bbagrm.2011.11.005

Callahan, B. J., McMurdie, P. J., & Holmes, S. P. (2017). Exact sequence variants should replace operational taxonomic units in marker-gene data analysis. ISME Journal, 11(12), 2639–2643. doi:10.1038/ismej.2017.119

Callahan, B. J., McMurdie, P. J., Rosen, M. J., Han, A. W., Johnson, A. J. A., & Holmes, S. P. (2016). DADA2: High-resolution sample inference from Illumina amplicon data. Nature Methods, 13(7), 581–583. doi:10.1038/nmeth.3869

Clare, E. L., Chain, F. J. J., Littlefair, J. E., & Cristescu, M. E. (2016). The effects of parameter choice on defining molecular operational taxonomic units and resulting ecological analyses of metabarcoding data. Genome, 59(11), 981–990. doi:10.1139/gen-2015-0184

Cock, P. J. A., Antao, T., Chang, J. T., Chapman, B. A., Cox, C. J., Dalke, A., … De Hoon, M. J. L. (2009). Biopython: Freely available Python tools for computational molecular biology and bioinformatics. Bioinformatics, 25(11), 1422–1423. doi:10.1093/bioinformatics/btp163

Creedy, T. J., Norman, H., Tang, C. Q., Qing Chin, K., Andujar, C., Arribas, P., … Vogler, A. P. (2019). A validated workflow for rapid taxonomic assignment and monitoring of a national fauna of bees (Apiformes) using high throughput DNA barcoding. Molecular Ecology Resources, 20(1), 40–53. doi:10.1111/1755-0998.13056

Delsuc, F., & Ranwez, V. (2020). Accurate alignment of (meta)barcoding data sets using MACSE. In C. Scornavacca, F. Delsuc, & N. Galtier (Eds.), Phylogenetics in the Genomic Era (hal-025411, pp. 2.3:1–2.3:31). No commercial publisher | Authors open access book.

Edgar, R. (2016). UNOISE2: improved error-correction for Illumina 16S and ITS amplicon sequencing. BioRxiv, 081257. doi:10.1101/081257

Edgar, R. C. (2013). UPARSE: highly accurate OTU sequences from microbial amplicon reads. Nature Methods, 10(10), 996–8. doi:10.1038/nmeth.2604

Edgar, R. C., Haas, B. J., Clemente, J. C., Quince, C., & Knight, R. (2011). UCHIME improves sensitivity and speed of chimera detection. Bioinformatics (Oxford, England), 27(16), 2194–200. doi:10.1093/bioinformatics/btr381

Elbrecht, V., Braukmann, T. W. A., Ivanova, N. V., Prosser, S. W. J., Hajibabaei, M., Wright, M., … Steinke, D. (2019). Validation of COI metabarcoding primers for terrestrial arthropods. PeerJ, 7:e7745. doi:http://doi.org/10.7717/peerj.7745

Elbrecht, V., Vamos, E. E., Steinke, D., & Leese, F. (2018). Estimating intraspecific genetic diversity from community DNA metabarcoding data. PeerJ, 6:e4644. doi:10.7287/peerj.preprints.3269v3

Flynn, J. M., Brown, E. a., Chain, F. J. J., MacIsaac, H. J., & Cristescu, M. E. (2015). Toward accurate molecular identification of species in complex environmental samples: testing the performance of sequence filtering and clustering methods. Ecology and Evolution, 5(11), 2252–2266. doi:10.1002/ece3.1497

Hamady, M., Walker, J. J., Harris, J. K., Gold, N. J., & Knight, R. (2008). Error-correcting barcoded primers for pyrosequencing hundreds of samples in multiplex. Nature Methods, 5(3), 235–237. doi:10.1038/nmeth.1184

Hazkani-Covo, E., Sorek, R., & Graur, D. (2003). Evolutionary dynamics of large Numts in the human genome: Rarity of independent insertions and abundance of post-insertion duplications. Journal of Molecular Evolution, 56(2), 169–174. doi:10.1007/s00239-002-2390-5

Huang, Y., Lu, W., Ji, J., Zhang, X., Zhang, P., & Chen, W. (2019). Heteroplasmy in the complete chicken mitochondrial genome. PLoS ONE, 14(11), 1–16. doi:10.1371/journal.pone.0224677

Huson, D. H., Beier, S., Flade, I., Górska, A., El-Hadidi, M., Mitra, S., … Tappu, R. (2016). MEGAN community edition - Interactive exploration and analysis of large-scale microbiome sequencing data. PLoS Computational Biology, 12(6), 1–12. doi:10.1371/journal.pcbi.1004957

Katoh, K., & Standley, D. M. (2013). MAFFT multiple sequence alignment software version 7: Improvements in performance and usability. Molecular Biology and Evolution, 30(4), 772–780. doi:10.1093/molbev/mst010

Liu, M., Clarke, L. J., Baker, S. C., Jordan, G. J., & Burridge, C. P. (2019). A practical guide to DNA metabarcoding for entomological ecologists. Ecological Entomology, 45, 373–385. doi:10.1111/een.12831

Lopez, J. V., Yuhki, N., Masuda, R., Modi, W., & O’Brien, S. J. (1994). Numt, a recent transfer and tandem amplification of mitochondrial DNA to the nuclear genome of the domestic cat. Journal of Molecular Evolution, 39(2), 174–190. doi:10.1007/BF00163806

Oliphant, T. E. (2006). Guide to NumPy. Methods (Vol. 1). USA: Trelgol Publishing. doi:10.1016/j.jmoldx.2015.02.001

Pamilo, P., Viljakainen, L., & Vihavainen, A. (2007). Exceptionally high density of NUMTs in the honeybee genome. Molecular Biology and Evolution, 24(6), 1340–1346. doi:10.1093/molbev/msm055

Paradis, E., Claude, J., & Strimmer, K. (2004). APE: Analyses of phylogenetics and evolution in R language. Bioinformatics, 20(2), 289–290. doi:10.1093/bioinformatics/btg412

Pons, J., & Vogler, A. P. (2005). Complex pattern of coalescence and fast evolution of a mitochondrial rRNA pseudogene in a recent radiation of tiger beetles. Molecular Biology and Evolution, 22(4), 991–1000. doi:10.1093/molbev/msi085

Quiros, P. M., Goyal, A., Jha, P., & Auwerx, J. (2017). Analysis of mtDNA/nDNA ratio in mice. Current Protocols in Mouse Biology, 7(1), 47. doi:10.1002/CPMO.21

Ramirez-Gonzalez, R., Yu, D. W., Bruce, C., Heavens, D., Caccamo, M., & Emerson, B. C. (2013). PyroClean: denoising pyrosequences from protein-coding amplicons for the recovery of interspecific and intraspecific genetic variation. PloS One, 8(3), e57615. doi:10.1371/journal.pone.0057615

Ramos, A., Barbena, E., Mateiu, L., del Mar González, M., Mairal, Q., Lima, M., … Santos, C. (2011). Nuclear insertions of mitochondrial origin: Database updating and usefulness in cancer studies. Mitochondrion, 11(6), 946–953. doi:10.1016/j.mito.2011.08.009

Rensch, T., Villar, D., Horvath, J., Odom, D. T., & Flicek, P. (2016). Mitochondrial heteroplasmy in vertebrates using ChIP-sequencing data. Genome Biology, 17(1), 1–14. doi:10.1186/s13059-016-0996-y

Richly, E., & Leister, D. (2004). NUMTs in sequenced eukaryotic genomes. Molecular Biology and Evolution, 21(6), 1081–1084. doi:10.1093/molbev/msh110

Schliep, K. P. (2011). phangorn: Phylogenetic analysis in R. Bioinformatics, 27(4), 592–593. doi:10.1093/bioinformatics/btq706

Schloss, P. D., & Westcott, S. L. (2011). Assessing and improving methods used in operational taxonomic unit-based approaches for 16S rRNA gene sequence analysis. Applied and Environmental Microbiology, 77(10), 3219–3226. doi:10.1128/AEM.02810-10

Shannon, P., Markiel, A., Ozier, O., Baliga, N. S., Wang, J. T., Ramage, D., … Ideker, T. (2003). Cytoscape: a software environment for integrated models of biomolecular interaction networks. Genome Research, 13(11), 2498–504. doi:10.1101/gr.1239303

Shi, H., Dong, J., Irwin, D. M., Zhang, S., & Mao, X. (2016). Repetitive transpositions of mitochondrial DNA sequences to the nucleus during the radiation of horseshoe bats (Rhinolophus, Chiroptera). Gene, 581(2), 161–169. doi:10.1016/j.gene.2016.01.035

Shokralla, S., Gibson, J. F., Nikbakht, H., Janzen, D. H., Hallwachs, W., & Hajibabaei, M. (2014). Next-generation DNA barcoding: Using next-generation sequencing to enhance and accelerate DNA barcode capture from single specimens. Molecular Ecology Resources, 14(5), 892–901. doi:10.1111/1755-0998.12236

Song, H., Buhay, J. E., Whiting, M. F., & Crandall, K. a. (2008). Many species in one: DNA barcoding overestimates the number of species when nuclear mitochondrial pseudogenes are coamplified. Proceedings of the National Academy of Sciences of the United States of America, 105(36), 13486–91. doi:10.1073/pnas.0803076105

Stamatakis, A. (2006). RAxML-VI-HPC: maximum likelihood-based phylogenetic analyses with thousands of taxa and mixed models. Bioinformatics (Oxford, England), 22(21), 2688–90. doi:10.1093/bioinformatics/btl446

Taberlet, P., Coissac, E., Pompanon, F., Brochmann, C., & Willerslev, E. (2012). Towards next-generation biodiversity assessment using DNA metabarcoding. Molecular Ecology, 21(8), 2045–50. doi:10.1111/j.1365-294X.2012.05470.x

Turon, X., Antich, A., Palacín, C., Præbel, K., & Wangensteen, O. S. (2019). From metabarcoding to metaphylogeography: separating the wheat from the chaff. BioRxiv, (May), 629535. doi:10.1101/629535

Wang, W. Y., Srivathsan, A., Foo, M., Yamane, S., & Meier, R. (2018). Sorting specimen-rich invertebrate samples with cost-effective NGS barcodes: validating a reverse workflow for specimen processing. Molecular Ecology Resources, 12(10), 3218–3221. doi:10.1111/1755-0998.12751

Yu, D., Ji, Y., Emerson, B., Wang, X., Ye, C., Yang, C., & Ding, Z. (2012). Biodiversity soup: metabarcoding of arthropods for rapid biodiversity assessment and biomonitoring. Methods in Ecology and Evolution, 3(4), 613–623. doi:10.1111/j.2041-210X.2012.00198.x

Zhang, J., Kapli, P., Pavlidis, P., & Stamatakis, A. (2013). A general species delimitation method with applications to phylogenetic placements. Bioinformatics (Oxford, England), 29(22), 2869–76. doi:10.1093/bioinformatics/btt499

