## SUPPLEMENTARY MATERIALS for "Validated removal of nuclear pseudogenes and sequencing artefacts from mitochondrial metabarcode data" (from another version of this preprint: version 1, 2020-06-18)

### for

**Summary:**

**1. Supplementary text for Material and Methods**

**1.1. Data generation:** Generation of the BEE dataset

**1.2. Derivation of formulae**

**1.3. Workflow of NUMTdumper script**

**2. Supplementary tables**

**2.1. Supplementary tables for the BEE Dataset (Tables S1-S5)**

**2.2. Supplementary tables for the COL Dataset (Tables S6-S10)**

**2.3. Supplementary tables for the CAC Dataset (Tables S11-S15)**

**3. Supplementary Figures**

**3.1. Schematic representation summarising the key steps in generating the amplicon sequence variant (ASV) datasets and NUMT dumping workflow (Figure S1)**

### 1. SUPPLEMENTARY TEXT FOR MATERIAL AND METHODS

#### 1.1. Data generation

Generation of the BEE dataset:

We selected 462 libraries (individuals) that fulfilled the following criteria: (i) the target barcode sequence (i.e., identified as the representative barcode sequence for the library) was a bee identified to species level against a UK regional bee barcode reference database (Creedy et al. 2019); (ii) the molecular ID of the target barcode matched the morphological ID of the specimen; and (iii) the library had between 1,000 and 40,000 reads. Fifty communities were generated by randomly selecting 100 libraries from among the 462 and sampling a random set of reads from each, subsequent to primer trimming, pair merging and quality filtering. The number of reads randomly selected from each library was a scaled function of the original number of reads from the library, to simulate a metabarcoding pool by retaining roughly equal numbers of reads per individual, with some variation. Denoising and chimera detection was finally applied to each mock community.

#### 1.2. Derivation of formulae

*Estimation of unknown counts*

We can estimate the total number of initial and filtered (surviving) target- and non-target ASVs based on (i) the initial and final numbers of ASVs and (ii) the proportion of retained target- and non-target ASVs, under the assumption that verified-target- and verified-non-target-ASVs are a representative subset of all target- and non-target-ASVs.

**Where we know:**

$T$= total initial ASVs
$t$= total filtered (surviving) ASVs
$A_v$= initial verified-authentic-ASVs
$a_v$= filtered (surviving) verified-authentic-ASVs
$N_v$= initial verified-non-authentic-ASVs
$n_v$= filtered (surviving) verified-non-authentic-ASVs
$paA_v$= proportion of surviving verified authentic-ASVs = $\frac{a_v}{A_v}$
$pnN_v$= proportion of surviving verified non-authentic-ASVs = $\frac{t_v}{T_v}$

**We want to estimate:**

$A$= total initial authentic-ASVs
$a$= total filtered (surviving) authentic-ASVs
$N$= total initial non-authentic-ASVs
$n$= total filtered (surviving) non-authentic-ASVs

**And we assume:**

$$paA\text{= proportion of surviving authentic-ASVs} = \frac{a}{A} \approx \frac{a_v}{A_v}$$

$$pnN\text{= proportion of surviving non-authentic-ASVs} = \frac{n}{N} \approx \frac{t_v}{T_v}$$

**Then, given:**

$$T = A + N \qquad t = a + n \qquad p_{aA} = \frac{a}{A} \qquad p_{nN} = \frac{n}{N}$$

$$\Rightarrow n = t - a \qquad \Rightarrow A = \frac{a}{p_{aA}} \qquad \Rightarrow N = \frac{n}{p_{nN}}$$

$$T = \frac{a}{p_{aA}} + \frac{n}{p_{nN}} = \frac{a}{p_{aA}} + \frac{t-a}{p_{nN}} = \frac{a}{p_{aA}} + \frac{t}{p_{nN}} - \frac{a}{p_{nN}} = \frac{1}{p_{nN}}t + a(\frac{1}{p_{aA}} - \frac{1}{p_{nN}})$$

$$\Rightarrow a(\frac{1}{p_{aA}} - \frac{1}{p_{nN}}) = T - \frac{1}{p_{nN}}t$$

$$\Rightarrow a = \frac{T - \frac{1}{p_{nN}}t}{\frac{1}{p_{aA}} - \frac{1}{p_{nN}}}$$

Assuming that $p_{aA} = \frac{a}{A} \approx \frac{a_v}{A_v}$ and $p_{nN} = \frac{n}{N} \approx \frac{n_v}{N_v}$:

$$a = \frac{\boldsymbol{T - \frac{N_v}{n_v}t}}{\boldsymbol{\frac{A_v}{a_v} - \frac{N_v}{n_v}}}$$

Equivalent derivation can be used to calculate $n$, $N$ and $A$. It should be noted that in theory this estimation is sensitive to the ratio of ASV survival proportions: highly uneven ratios, or values of 0, will not compute or will output very inaccurate values. However, in practice most real-world values of initial and surviving validated ASV counts will generate reasonable estimated ASV counts.

*Beta-development of an optimisation index (Not used in the main text)*

To automatically suggest one or more ideal sets of filtering thresholds, we calculate a standardised success score, $s$, with values between 0 and 1. We consider the ideal filtering outcome to be where a) the proportion of estimated surviving non-authentic-ASVs ($\frac{n}{t}$)in the dataset is minimised, while b) the proportion of estimated surviving authentic-ASVs relative to initial authentic-ASVs ($\frac{a}{A}$)is maximised. Thus,

$$s = 0.5 \times (1 - \frac{a}{A} + \frac{n}{t})$$

This computation assigns equal weight to NUMT removal and authentic ASV retention, however different applications may wish to optimise towards one or the other. Thus we can extend this score using a weighting value ($w$) between 0 (only interested in optimal rejection of non-authentic-ASVs) and 1 (only interested in optimal retention of authentic-ASVs), i.e.

$$s = 0.5 \times (w(1 - \frac{a}{A}) + (1 - w)\frac{n}{t})$$

The value of s ranges between 0 (optimal) and 1, and can be weighted towards rejection of na-ASVs by using values of w greater than 0.5 and towards retention of a-ASVs with values of w less than 0.5. Furthermore, *NUMTdumper* can perform this optimisation either on an unlimited basis, or within defined limits of na-ASV rejection and/or a-ASV retention, depending on the use case - for example, finding the optimal threshold set for a highly stringent filter where no more than 5% of the final ASVs are estimated to be non-authentic, or for a conservative filter where at least 95% of estimated authentic ASVs are retained.

##### 1.3. Workflow of *NUMTdumper* script

The *NUMTdumper* script is comprised of two key parts, assessment, and self-validation (Fig. 1C of main text). During assessment, the abundance distributions of all input ASVs are interrogated within sample, clade and/or taxon, and ASVs survival is established for different filtering strategies and abundance thresholds. Self-validation comprises firstly the separate assignment of input ASVs as validated-authentic-ASVs (va-ASVs), validated-non-authentic-ASVs (vna-ASVs), or unclassified-ASVs (u-ASVs), as described above. Secondly, *NUMTdumper* evaluates (i) the removal efficiency of NUMTs and other erroneous sequences (non-authentic-ASVs; na-ASVs) and (ii) the collateral effect on the removal of authentic haplotypes (authentic-ASVs; a-ASVs), as a function of the different filtering strategies and read abundance thresholds. *NUMTdumper* facilitates the identification of filtering parameters that optimise the removal of na-ASVs as a function of a-ASVs retention, providing approximate measures of prevalence of both a-ASVs and na-ASVs in the final dataset.

*NUMTdumper* requires as inputs (i) the ASV sequences, (ii) the set of reads for each metabarcoding library, and (iii) an external reference set of sequences expected to be present among the reads of the focal libraries to assign va-ASVs. The assignment as vna-ASVs is based on the presence of frameshift and stop mutations. In addition, (iv) taxonomic identity of the ASVs could be provided for applying read abundance filtering within taxonomic classes (here done only for the CAC dataset). The workflow comprises the following steps:

**Assessment I: Binning and counting**. Within a dataset, reads for each library are queried against the ASVs using exact matches to identify ASV incidence and count the number of each ASV present in each library. The total number of reads for each ASV across all libraries is also calculated.

To identify clades, a UPGMA tree of the ASVs is constructed using F84 model-corrected distances (Felsenstein & Churchill, 1996) based on a MAFFT FFT-NS-2 alignment (Katoh et al. 2002) of the ASV sequences. Clades are delimited based on a specified divergence threshold, with each ASV assigned to a clade. As an alternative, *NUMTdumper* also allows the definition of clade assignation based on a provided file with the taxonomic identity of the ASVs, which allows for the use of filtering criteria within custom defined taxonomic ranks (e.g., when multiple phyla are present in the dataset, each ASV could be assigned to a phylum). These two counts (ASVs in a library and total ASVs) and two bins (clade and taxonomic rank) permit the computation of counts of read numbers for all ASVs for any combination of count and bin(s).

**Assessment II: Filtering.** Within a dataset, the absolute and/or relative frequency of each ASV is assessed for each specified count/bin criterion against a specified set of thresholds, and where an ASV count/proportion does not meet a minimum threshold in all bins in which it occurs, it is removed under that particular threshold; otherwise, it is retained.

**Self-Validation**. Within a dataset, ASVs are firstly designated as va-ASVs, vna-ASVs, or u-ASVs based on a perfect match to the dataset's reference set using BLASTn (va-ASV), the presence of indels and/or stop codons in the translation (vna-ASV), or neither of these (u-ASV). Then, for each binning criterion and threshold, the rejection or retention of each va-ASV or vna-ASV is recorded. These data are used to compute, for each threshold, (i) the proportion of false negatives i.e., the proportion of va-ASVs that are removed, and (ii) the proportion of false positives i.e., the proportion of vna-ASVs that are retained. This analysis is based on the fact that a subset (the va-ASVs) of the authentic mitochondrial ASVs (a-ASVs) are known, and their presence or absence ('survival') after the filtering process can be established. Survival can be assessed equally for the copies that are known to be NUMTs or erroneous based on their frameshifts or stop codons (vna-ASV). To evaluate parameter optimality to minimise va-ASV removal and vna-ASV survival, trends of the proportion of false negatives and false positives were estimated and plotted for every threshold within each of the five filtering criteria described above.

### 2. SUPPLEMENTARY TABLES

#### 2.1. Supplementary tables for the BEE Dataset (Tables S1-S5)

**Table S1.** Filtering performance of filtering criteria "Absolute ASV abundance by library" for a range of minimum threshold values for read abundance for the BEE dataset.

| Criterium | Thershold | Surviving ASVs (t)* | Surviving va-ASVs (av)* | Surviving vna-ASVs (nv)* | Initial a-ASVs (A)** | Surviving a-ASV (a)** | Initial na-ASVs (N)** | Surviving na-ASV (n)** | n/t *** |
|---|---|---|---|---|---|---|---|---|---|
| **Pre-filtering initial values** | | **T = 2251** | **Av = 160** | **Nv = 117** | | | | | |
| **Absolute ASV abundance by library** (Threshold = minimum threshold of read-abundance in a given library) | 3 | 1128 (50.1%) | 160 (100%) | 62 (53%) | -137.9 | -137.9 | 2388.9 | 1265.9 | 1.12 |
| | 4 | 797 (35.4%) | 160 (100%) | 35 (29.9%) | 176.4 | 176.4 | 2074.6 | 620.6 | 0.78 |
| | 5 | 626 (27.8%) | 160 (100%) | 26 (22.2%) | 161.7 | 161.7 | 2089.3 | 464.3 | 0.74 |
| | 8 | 400 (17.8%) | 159 (99.4%) | 14 (12%) | 149.5 | 148.5 | 2101.5 | 251.5 | 0.63 |
| | 10 | 345 (15.3%) | 159 (99.4%) | 10 (8.5%) | 168.0 | 167.0 | 2083.0 | 178.0 | 0.52 |
| | 12 | 309 (13.7%) | 159 (99.4%) | 9 (7.7%) | 148.2 | 147.2 | 2102.8 | 161.8 | 0.52 |
| | 15 | 279 (12.4%) | 159 (99.4%) | 7 (6%) | 154.5 | 153.6 | 2096.5 | 125.4 | 0.45 |
| | 20 | 249 (11.1%) | 159 (99.4%) | 5 (4.3%) | 160.7 | 159.7 | 2090.3 | 89.3 | 0.36 |
| | 25 | 225 (10%) | 156 (97.5%) | 3 (2.6%) | 176.2 | 171.8 | 2074.8 | 53.2 | 0.24 |
| | 35 | 194 (8.6%) | 152 (95%) | 1 (0.9%) | 185.6 | 176.3 | 2065.4 | 17.7 | 0.09 |
| | 40 | 182 (8.1%) | 149 (93.1%) | 1 (0.9%) | 176.4 | 164.3 | 2074.6 | 17.7 | 0.10 |
| | 45 | 177 (7.9%) | 148 (92.5%) | 1 (0.9%) | 172.1 | 159.2 | 2078.9 | 17.8 | 0.10 |
| | 50 | 172 (7.6%) | 146 (91.3%) | 1 (0.9%) | 169.0 | 154.2 | 2082.0 | 17.8 | 0.10 |
| | 60 | 165 (7.3%) | 143 (89.4%) | 1 (0.9%) | 164.7 | 147.2 | 2086.3 | 17.8 | 0.11 |
| | 70 | 154 (6.8%) | 142 (88.8%) | 0 (0%) | 173.5 | 154.0 | 2077.5 | 0.0 | 0.00 |
| | 80 | 151 (6.7%) | 140 (87.5%) | 0 (0%) | 172.6 | 151.0 | 2078.4 | 0.0 | 0.00 |
| | 90 | 146 (6.5%) | 137 (85.6%) | 0 (0%) | 170.5 | 146.0 | 2080.5 | 0.0 | 0.00 |
| | 100 | 143 (6.4%) | 136 (85%) | 0 (0%) | 168.2 | 143.0 | 2082.8 | 0.0 | 0.00 |

* Observed values; ** Estimated values; *** The ratio n/t represents the estimated proportion of na-ASVs in the filtered dataset.

**Table S2.** Filtering performance of filtering criteria "Relative ASV abundance by library" for a range of minimum threshold values for read abundance for the BEE dataset.

| Criterium | Thershold | Surviving ASVs (t)* | Surviving va-ASVs (av)* | Surviving vna-ASVs (nv)* | Initial a-ASVs (A)** | Surviving a-ASV (a)** | Initial na-ASVs (N)** | Surviving na-ASV (n)** | n/t *** |
|---|---|---|---|---|---|---|---|---|---|
| **Pre-filtering initial values** | | T = 2251 | Av = 160 | Nv = 117 | | | | | |
| **Relative ASV abundance by library** (Threshold = minimum proportion of read-abundance relative to the total number of reads in that library) | 0.00025 | 393 (17.5%) | 159 (99.4%) | 14 (12%) | 141.5 | 140.6 | 2109.5 | 252.4 | 0.64 |
| | 0.0005 | 270 (12%) | 159 (99.4%) | 7 (6%) | 144.9 | 144.0 | 2106.1 | 126.0 | 0.47 |
| | 0.001 | 206 (9.2%) | 155 (96.9%) | 1 (0.9%) | 194.5 | 188.4 | 2056.5 | 17.6 | 0.09 |
| | 0.0015 | 173 (7.7%) | 147 (91.9%) | 1 (0.9%) | 168.9 | 155.2 | 2082.1 | 17.8 | 0.10 |
| | 0.002 | 163 (7.2%) | 143 (89.4%) | 1 (0.9%) | 162.4 | 145.1 | 2088.6 | 17.9 | 0.11 |
| | 0.0025 | 150 (6.7%) | 140 (87.5%) | 0 (0%) | 171.4 | 150.0 | 2079.6 | 0.0 | 0.00 |
| | 0.003 | 144 (6.4%) | 136 (85%) | 0 (0%) | 169.4 | 144.0 | 2081.6 | 0.0 | 0.00 |
| | 0.0035 | 142 (6.3%) | 135 (84.4%) | 0 (0%) | 168.3 | 142.0 | 2082.7 | 0.0 | 0.00 |
| | 0.004 | 142 (6.3%) | 135 (84.4%) | 0 (0%) | 168.3 | 142.0 | 2082.7 | 0.0 | 0.00 |
| | 0.0045 | 136 (6%) | 131 (81.9%) | 0 (0%) | 166.1 | 136.0 | 2084.9 | 0.0 | 0.00 |
| | 0.005 | 132 (5.9%) | 129 (80.6%) | 0 (0%) | 163.7 | 132.0 | 2087.3 | 0.0 | 0.00 |
| | 0.0055 | 128 (5.7%) | 126 (78.8%) | 0 (0%) | 162.5 | 128.0 | 2088.5 | 0.0 | 0.00 |
| | 0.006 | 121 (5.4%) | 120 (75%) | 0 (0%) | 161.3 | 121.0 | 2089.7 | 0.0 | 0.00 |
| | 0.0065 | 116 (5.2%) | 115 (71.9%) | 0 (0%) | 161.4 | 116.0 | 2089.6 | 0.0 | 0.00 |
| | 0.007 | 108 (4.8%) | 107 (66.9%) | 0 (0%) | 161.5 | 108.0 | 2089.5 | 0.0 | 0.00 |
| | 0.008 | 103 (4.6%) | 103 (64.4%) | 0 (0%) | 160.0 | 103.0 | 2091.0 | 0.0 | 0.00 |
| | 0.009 | 95 (4.2%) | 95 (59.4%) | 0 (0%) | 160.0 | 95.0 | 2091.0 | 0.0 | 0.00 |
| | 0.01 | 91 (4%) | 91 (56.9%) | 0 (0%) | 160.0 | 91.0 | 2091.0 | 0.0 | 0.00 |

* Observed values; ** Estimated values; *** The ratio n/t represents the estimated proportion of na-ASVs in the filtered dataset

**Table S3.** Filtering performance of filtering criteria "Relative ASV abundance by library and 15% clade" for a range of minimum threshold values for read abundance for the BEE dataset.

| Criterium | Thershold | Surviving ASVs (t)* | Surviving va-ASVs (av)* | Surviving vna-ASVs (nv)* | Initial a-ASVs (A)** | Surviving a-ASV (a)** | Initial na-ASVs (N)** | Surviving na-ASV (n)** | n/t *** |
|---|---|---|---|---|---|---|---|---|---|
| **Pre-filtering initial values** | | T = 2251 | Av = 160 | Nv = 117 | | | | | |
| **Relative ASV abundance by library and 15% clade** (Threshold = minimum read proportion relative to the total number of reads in the 15% divergence clade where each ASV is included in that library) | 0.001 | 1574 (69.9%) | 160 (100%) | 88 (75.2%) | -480.3 | -480.3 | 2731.3 | 2054.3 | 1.31 |
| | 0.0025 | 1149 (51%) | 160 (100%) | 68 (58.1%) | -380.3 | -380.3 | 2631.3 | 1529.3 | 1.33 |
| | 0.005 | 893 (39.7%) | 160 (100%) | 52 (44.4%) | -193.4 | -193.4 | 2444.4 | 1086.4 | 1.22 |
| | 0.01 | 621 (27.6%) | 160 (100%) | 35 (29.9%) | -74.7 | -74.7 | 2325.7 | 695.7 | 1.12 |
| | 0.015 | 491 (21.8%) | 159 (99.4%) | 25 (21.4%) | 12.8 | 12.8 | 2238.2 | 478.2 | 0.97 |
| | 0.02 | 419 (18.6%) | 159 (99.4%) | 22 (18.8%) | -5.3 | -5.3 | 2256.3 | 424.3 | 1.01 |
| | 0.025 | 368 (16.3%) | 158 (98.8%) | 21 (17.9%) | -44.6 | -44.0 | 2295.6 | 412.0 | 1.12 |
| | 0.03 | 331 (14.7%) | 157 (98.1%) | 18 (15.4%) | -18.5 | -18.2 | 2269.5 | 349.2 | 1.05 |
| | 0.035 | 316 (14%) | 157 (98.1%) | 17 (14.5%) | -13.2 | -13.0 | 2264.2 | 329.0 | 1.04 |
| | 0.04 | 296 (13.1%) | 157 (98.1%) | 15 (12.8%) | 8.7 | 8.5 | 2242.3 | 287.5 | 0.97 |
| | 0.045 | 287 (12.7%) | 157 (98.1%) | 15 (12.8%) | -1.9 | -1.8 | 2252.9 | 288.8 | 1.01 |
| | 0.05 | 268 (11.9%) | 157 (98.1%) | 14 (12%) | -1.6 | -1.5 | 2252.6 | 269.5 | 1.01 |
| | 0.055 | 263 (11.7%) | 156 (97.5%) | 12 (10.3%) | 36.8 | 35.9 | 2214.2 | 227.1 | 0.86 |
| | 0.06 | 258 (11.5%) | 156 (97.5%) | 12 (10.3%) | 31.1 | 30.3 | 2219.9 | 227.7 | 0.88 |
| | 0.065 | 248 (11%) | 156 (97.5%) | 11 (9.4%) | 41.3 | 40.2 | 2209.7 | 207.8 | 0.84 |
| | 0.075 | 236 (10.5%) | 155 (96.9%) | 10 (8.5%) | 49.4 | 47.8 | 2201.6 | 188.2 | 0.80 |
| | 0.09 | 219 (9.7%) | 152 (95%) | 10 (8.5%) | 30.8 | 29.2 | 2220.2 | 189.8 | 0.87 |
| | 0.1 | 213 (9.5%) | 149 (93.1%) | 10 (8.5%) | 24.4 | 22.7 | 2226.6 | 190.3 | 0.89 |
| | 0.15 | 184 (8.2%) | 137 (85.6%) | 8 (6.8%) | 38.2 | 32.7 | 2212.8 | 151.3 | 0.82 |
| | 0.2 | 170 (7.6%) | 134 (83.8%) | 8 (6.8%) | 20.9 | 17.5 | 2230.1 | 152.5 | 0.90 |
| | 0.25 | 157 (7%) | 126 (78.8%) | 7 (6%) | 30.7 | 24.2 | 2220.3 | 132.8 | 0.85 |
| | 0.3 | 143 (6.4%) | 117 (73.1%) | 7 (6%) | 12.4 | 9.1 | 2238.6 | 133.9 | 0.94 |
| | 0.35 | 137 (6.1%) | 113 (70.6%) | 7 (6%) | 3.6 | 2.5 | 2247.4 | 134.5 | 0.98 |
| | 0.4 | 133 (5.9%) | 110 (68.8%) | 6 (5.1%) | 27.6 | 19.0 | 2223.4 | 114.0 | 0.86 |
| | 0.5 | 118 (5.2%) | 96 (60%) | 5 (4.3%) | 39.1 | 23.5 | 2211.9 | 94.5 | 0.80 |
| | 0.6 | 110 (4.9%) | 91 (56.9%) | 4 (3.4%) | 61.8 | 35.2 | 2189.2 | 74.8 | 0.68 |
| | 0.7 | 102 (4.5%) | 85 (53.1%) | 4 (3.4%) | 50.4 | 26.8 | 2200.6 | 75.2 | 0.74 |
| | 0.8 | 96 (4.3%) | 79 (49.4%) | 4 (3.4%) | 41.4 | 20.5 | 2209.6 | 75.5 | 0.79 |
| | 0.9 | 90 (4%) | 73 (45.6%) | 4 (3.4%) | 30.9 | 14.1 | 2220.1 | 75.9 | 0.84 |

* Observed values; ** Estimated values; *** The ratio n/t represents the estimated proportion of na-ASVs in the filtered dataset

**Table S4.** Filtering performance of filtering criteria “Relative ASV abundance by library and 20% clade” for a range of minimum threshold values for read abundance for the BEE dataset.

| Criterium | Thershold | Surviving ASVs (t)* | Surviving va-ASVs (av)* | Surviving vna-ASVs (nv)* | Initial a-ASVs (A)** | Surviving a-ASV (a)** | Initial na-ASVs (N)** | Surviving na-ASV (n)** | n/t *** |
|---|---|---|---|---|---|---|---|---|---|
| **Pre-filtering initial values** | | T = 2251 | Av = 160 | Nv = 117 | | | | | |
| **Relative ASV abundance by library and 20% clade** (Threshold = minimum read proportion relative to the total number of reads in the 20% divergence clade where each ASV is included in that library) | 0.001 | 776 (34.5%) | 160 (100%) | 53 (45.3%) | -445.5 | -445.5 | 2696.5 | 1221.5 | 1.57 |
| | 0.0025 | 626 (27.8%) | 160 (100%) | 47 (40.2%) | -465.1 | -465.1 | 2716.1 | 1091.1 | 1.74 |
| | 0.005 | 488 (21.7%) | 158 (98.8%) | 33 (28.2%) | -208.2 | -205.6 | 2459.2 | 693.6 | 1.42 |
| | 0.01 | 344 (15.3%) | 155 (96.9%) | 18 (15.4%) | -2.8 | -2.7 | 2253.8 | 346.7 | 1.01 |
| | 0.015 | 288 (12.8%) | 151 (94.4%) | 14 (12%) | 22.6 | 21.4 | 2228.4 | 266.6 | 0.93 |
| | 0.02 | 249 (11.1%) | 146 (91.3%) | 12 (10.3%) | 22.4 | 20.4 | 2228.6 | 228.6 | 0.92 |
| | 0.025 | 226 (10%) | 142 (88.8%) | 11 (9.4%) | 18.1 | 16.1 | 2232.9 | 209.9 | 0.93 |
| | 0.03 | 204 (9.1%) | 134 (83.8%) | 11 (9.4%) | -10.3 | -8.6 | 2261.3 | 212.6 | 1.04 |
| | 0.035 | 186 (8.3%) | 127 (79.4%) | 9 (7.7%) | 17.9 | 14.2 | 2233.1 | 171.8 | 0.92 |
| | 0.04 | 176 (7.8%) | 124 (77.5%) | 9 (7.7%) | 4.1 | 3.2 | 2246.9 | 172.8 | 0.98 |
| | 0.045 | 168 (7.5%) | 118 (73.8%) | 9 (7.7%) | -7.8 | -5.8 | 2258.8 | 173.8 | 1.03 |
| | 0.05 | 155 (6.9%) | 116 (72.5%) | 8 (6.8%) | 1.7 | 1.2 | 2249.3 | 153.8 | 0.99 |
| | 0.055 | 148 (6.6%) | 111 (69.4%) | 8 (6.8%) | -9.5 | -6.6 | 2260.5 | 154.6 | 1.04 |
| | 0.06 | 139 (6.2%) | 105 (65.6%) | 8 (6.8%) | -25.4 | -16.6 | 2276.4 | 155.6 | 1.12 |
| | 0.065 | 134 (6%) | 103 (64.4%) | 8 (6.8%) | -34.6 | -22.3 | 2285.6 | 156.3 | 1.17 |
| | 0.075 | 127 (5.6%) | 98 (61.3%) | 8 (6.8%) | -49.5 | -30.3 | 2300.5 | 157.3 | 1.24 |
| | 0.09 | 112 (5%) | 87 (54.4%) | 8 (6.8%) | -88.2 | -47.9 | 2339.2 | 159.9 | 1.43 |
| | 0.1 | 103 (4.6%) | 81 (50.6%) | 8 (6.8%) | -116.3 | -58.9 | 2367.3 | 161.9 | 1.57 |
| | 0.15 | 81 (3.6%) | 65 (40.6%) | 6 (5.1%) | -97.0 | -39.4 | 2348.0 | 120.4 | 1.49 |
| | 0.2 | 66 (2.9%) | 53 (33.1%) | 6 (5.1%) | -176.6 | -58.5 | 2427.6 | 124.5 | 1.89 |
| | 0.25 | 59 (2.6%) | 50 (31.3%) | 5 (4.3%) | -137.9 | -43.1 | 2388.9 | 102.1 | 1.73 |
| | 0.3 | 56 (2.5%) | 48 (30%) | 5 (4.3%) | -156.2 | -46.9 | 2407.2 | 102.9 | 1.84 |
| | 0.35 | 53 (2.4%) | 45 (28.1%) | 5 (4.3%) | -181.1 | -50.9 | 2432.1 | 103.9 | 1.96 |
| | 0.4 | 52 (2.3%) | 44 (27.5%) | 5 (4.3%) | -190.3 | -52.3 | 2441.3 | 104.3 | 2.01 |
| | 0.5 | 48 (2.1%) | 40 (25%) | 5 (4.3%) | -232.5 | -58.1 | 2483.5 | 106.1 | 2.21 |
| | 0.6 | 43 (1.9%) | 37 (23.1%) | 4 (3.4%) | -172.3 | -39.8 | 2423.3 | 82.8 | 1.93 |
| | 0.7 | 40 (1.8%) | 34 (21.3%) | 4 (3.4%) | -207.3 | -44.0 | 2458.3 | 84.0 | 2.10 |
| | 0.8 | 36 (1.6%) | 30 (18.8%) | 4 (3.4%) | -267.1 | -50.1 | 2518.1 | 86.1 | 2.39 |
| | 0.9 | 30 (1.3%) | 24 (15%) | 4 (3.4%) | -405.5 | -60.8 | 2656.5 | 90.8 | 3.03 |

* Observed values; ** Estimated values; *** The ratio n/t represents the estimated proportion of na-ASVs in the filtered dataset

**Table S5.** Filtering performance of filtering criteria "Relative ASV abundance by library and 26% clade" for a range of minimum threshold values for read abundance for the BEE dataset.

| Criterium | Thershold | Surviving ASVs (t)* | Surviving va-ASVs (av)* | Surviving vna-ASVs (nv)* | Initial a-ASVs (A)** | Surviving a-ASV (a)** | Initial na-ASVs (N)** | Surviving na-ASV (n)** | n/t *** |
|---|---|---|---|---|---|---|---|---|---|
| **Pre-filtering initial values** | | T = 2251 | Av = 160 | Nv = 117 | | | | | |
| **Relative ASV abundance by library and 26% clade** (Threshold = minimum read proportion relative to the total number of reads in the 26% divergence clade where each ASV is included in that library) | 0.001 | 490 (21.8%) | 159 (99.4%) | 22 (18.8%) | 82.8 | 82.3 | 2168.2 | 407.7 | 0.83 |
| | 0.0025 | 374 (16.6%) | 158 (98.8%) | 16 (13.7%) | 77.8 | 76.8 | 2173.2 | 297.2 | 0.79 |
| | 0.005 | 295 (13.1%) | 153 (95.6%) | 12 (10.3%) | 75.1 | 71.8 | 2175.9 | 223.2 | 0.76 |
| | 0.01 | 205 (9.1%) | 141 (88.1%) | 6 (5.1%) | 107.9 | 95.1 | 2143.1 | 109.9 | 0.54 |
| | 0.015 | 179 (8%) | 135 (84.4%) | 5 (4.3%) | 103.4 | 87.2 | 2147.6 | 91.8 | 0.51 |
| | 0.02 | 155 (6.9%) | 127 (79.4%) | 4 (3.4%) | 102.7 | 81.6 | 2148.3 | 73.4 | 0.47 |
| | 0.025 | 143 (6.4%) | 118 (73.8%) | 4 (3.4%) | 93.9 | 69.3 | 2157.1 | 73.7 | 0.52 |
| | 0.03 | 135 (6%) | 113 (70.6%) | 4 (3.4%) | 86.4 | 61.0 | 2164.6 | 74.0 | 0.55 |
| | 0.035 | 118 (5.2%) | 104 (65%) | 3 (2.6%) | 96.6 | 62.8 | 2154.4 | 55.2 | 0.47 |
| | 0.04 | 108 (4.8%) | 95 (59.4%) | 3 (2.6%) | 88.5 | 52.6 | 2162.5 | 55.4 | 0.51 |
| | 0.045 | 102 (4.5%) | 90 (56.3%) | 3 (2.6%) | 82.5 | 46.4 | 2168.5 | 55.6 | 0.55 |
| | 0.05 | 100 (4.4%) | 88 (55%) | 3 (2.6%) | 80.6 | 44.3 | 2170.4 | 55.7 | 0.56 |
| | 0.055 | 88 (3.9%) | 77 (48.1%) | 3 (2.6%) | 66.5 | 32.0 | 2184.5 | 56.0 | 0.64 |
| | 0.06 | 79 (3.5%) | 69 (43.1%) | 3 (2.6%) | 52.5 | 22.6 | 2198.5 | 56.4 | 0.71 |
| | 0.065 | 78 (3.5%) | 68 (42.5%) | 3 (2.6%) | 50.8 | 21.6 | 2200.2 | 56.4 | 0.72 |
| | 0.075 | 69 (3.1%) | 60 (37.5%) | 3 (2.6%) | 32.3 | 12.1 | 2218.7 | 56.9 | 0.82 |
| | 0.09 | 54 (2.4%) | 47 (29.4%) | 3 (2.6%) | -13.9 | -4.1 | 2264.9 | 58.1 | 1.08 |
| | 0.1 | 48 (2.1%) | 41 (25.6%) | 3 (2.6%) | -42.1 | -10.8 | 2293.1 | 58.8 | 1.22 |
| | 0.15 | 31 (1.4%) | 25 (15.6%) | 3 (2.6%) | -204.6 | -32.0 | 2455.6 | 63.0 | 2.03 |
| | 0.2 | 20 (0.9%) | 16 (10%) | 3 (2.6%) | -507.2 | -50.7 | 2758.2 | 70.7 | 3.54 |
| | 0.25 | 17 (0.8%) | 14 (8.8%) | 2 (1.7%) | -305.1 | -26.7 | 2556.1 | 43.7 | 2.57 |
| | 0.3 | 16 (0.7%) | 13 (8.1%) | 2 (1.7%) | -350.4 | -28.5 | 2601.4 | 44.5 | 2.78 |
| | 0.35 | 13 (0.6%) | 11 (6.9%) | 2 (1.7%) | -493.2 | -33.9 | 2744.2 | 46.9 | 3.61 |
| | 0.4 | 12 (0.5%) | 10 (6.3%) | 2 (1.7%) | -583.2 | -36.4 | 2834.2 | 48.4 | 4.04 |
| | 0.5 | 11 (0.5%) | 9 (5.6%) | 2 (1.7%) | -701.8 | -39.5 | 2952.8 | 50.5 | 4.59 |
| | 0.6 | 10 (0.4%) | 9 (5.6%) | 1 (0.9%) | -193.7 | -10.9 | 2444.7 | 20.9 | 2.09 |
| | 0.7 | 5 (0.2%) | 4 (2.5%) | 1 (0.9%) | -865.5 | -21.6 | 3116.5 | 26.6 | 5.33 |
| | 0.8 | 3 (0.1%) | 2 (1.3%) | 1 (0.9%) | -4108.1 | -51.4 | 6359.1 | 54.4 | 18.12 |
| | 0.9 | 2 (0.1%) | 1 (0.6%) | 1 (0.9%) | 7505.1 | 46.9 | -5254.1 | -44.9 | -22.45 |

* Observed values; ** Estimated values; *** The ratio n/t represents the estimated proportion of na-ASVs in the filtered dataset

#### 2.2. Supplementary tables for the COL Dataset (Tables S6-S10)

**Table S6.** Filtering performance of filtering criteria "Absolute ASV abundance by library" for a range of minimum threshold values for read abundance for the COL dataset.

| Criterium | Thershold | Surviving ASVs (t)* | Surviving va-ASVs (av)* | Surviving vna-ASVs (nv)* | Initial a-ASVs (A)** | Surviving a-ASV (a)** | Initial na-ASVs (N)** | Surviving na-ASV (n)** | n/t *** |
|---|---|---|---|---|---|---|---|---|---|
| **Pre-filtering initial values** | | **T = 1845** | **Av = 74** | **Nv = 228** | | | | | |
| **Absolute ASV abundance by library** (Threshold = minimum threshold of read-abundance in a given library) | 3 | 1350 (73.2%) | 74 (100%) | 143 (62.7%) | 517.2 | 517.2 | 1327.8 | 832.8 | 0.617 |
| | 4 | 1139 (61.7%) | 74 (100%) | 105 (46.1%) | 536.3 | 536.3 | 1308.7 | 602.7 | 0.529 |
| | 5 | 1008 (54.6%) | 74 (100%) | 76 (33.3%) | 589.5 | 589.5 | 1255.5 | 418.5 | 0.415 |
| | 8 | 787 (42.7%) | 68 (91.9%) | 41 (18%) | 615.9 | 566.0 | 1229.1 | 221.0 | 0.281 |
| | 10 | 706 (38.3%) | 67 (90.5%) | 32 (14%) | 584.3 | 529.1 | 1260.7 | 176.9 | 0.251 |
| | 12 | 662 (35.9%) | 65 (87.8%) | 24 (10.5%) | 605.1 | 531.5 | 1239.9 | 130.5 | 0.197 |
| | 15 | 598 (32.4%) | 64 (86.5%) | 18 (7.9%) | 575.6 | 497.8 | 1269.4 | 100.2 | 0.168 |
| | 20 | 526 (28.5%) | 57 (77%) | 10 (4.4%) | 612.7 | 472.0 | 1232.3 | 54.0 | 0.103 |
| | 25 | 482 (26.1%) | 57 (77%) | 5 (2.2%) | 590.0 | 454.5 | 1255.0 | 27.5 | 0.057 |
| | 35 | 413 (22.4%) | 51 (68.9%) | 2 (0.9%) | 583.2 | 401.9 | 1261.8 | 11.1 | 0.027 |
| | 40 | 400 (21.7%) | 51 (68.9%) | 1 (0.4%) | 572.3 | 394.4 | 1272.7 | 5.6 | 0.014 |
| | 45 | 371 (20.1%) | 48 (64.9%) | 0 (0%) | 572.0 | 371.0 | 1273.0 | 0.0 | 0.000 |
| | 50 | 355 (19.2%) | 44 (59.5%) | 0 (0%) | 597.0 | 355.0 | 1248.0 | 0.0 | 0.000 |
| | 60 | 336 (18.2%) | 40 (54.1%) | 0 (0%) | 621.6 | 336.0 | 1223.4 | 0.0 | 0.000 |
| | 70 | 313 (17%) | 33 (44.6%) | 0 (0%) | 701.9 | 313.0 | 1143.1 | 0.0 | 0.000 |
| | 80 | 299 (16.2%) | 38 (51.4%) | 0 (0%) | 582.3 | 299.0 | 1262.7 | 0.0 | 0.000 |
| | 90 | 280 (15.2%) | 35 (47.3%) | 0 (0%) | 592.0 | 280.0 | 1253.0 | 0.0 | 0.000 |
| | 100 | 262 (14.2%) | 34 (45.9%) | 0 (0%) | 570.2 | 262.0 | 1274.8 | 0.0 | 0.000 |

* Observed values; ** Estimated values; *** The ratio n/t represents the estimated proportion of na-ASVs in the filtered dataset.

**Table S7.** Filtering performance of filtering criteria "Relative ASV abundance by library" for a range of minimum threshold values for read abundance for the COL dataset.

| Criterium | Thershold | Surviving ASVs (t)* | Surviving va-ASVs (av)* | Surviving vna-ASVs (nv)* | Initial a-ASVs (A)** | Surviving a-ASV (a)** | Initial na-ASVs (N)** | Surviving na-ASV (n)** | n/t *** |
|---|---|---|---|---|---|---|---|---|---|
| **Pre-filtering initial values** | | T = 1845 | Av = 74 | Nv = 228 | | | | | |
| | 0.00025 | 1813 (98.3%) | 74 (100%) | 225 (98.7%) | -587.0 | -587.0 | 2432.0 | 2400.0 | 1.324 |
| | 5.00E-04 | 1666 (90.3%) | 74 (100%) | 201 (88.2%) | 333.4 | 333.4 | 1511.6 | 1332.6 | 0.800 |
| | 0.00075 | 1499 (81.2%) | 74 (100%) | 170 (74.6%) | 484.9 | 484.9 | 1360.1 | 1014.1 | 0.677 |
| | 0.001 | 1381 (74.9%) | 74 (100%) | 147 (64.5%) | 538.9 | 538.9 | 1306.1 | 842.1 | 0.610 |
| | 0.0015 | 1159 (62.8%) | 74 (100%) | 106 (46.5%) | 563.0 | 563.0 | 1282.0 | 596.0 | 0.514 |
| | 0.002 | 1026 (55.6%) | 74 (100%) | 84 (36.8%) | 548.3 | 548.3 | 1296.8 | 477.8 | 0.466 |
| | 0.0025 | 920 (49.9%) | 72 (97.3%) | 65 (28.5%) | 572.8 | 557.3 | 1272.2 | 362.7 | 0.394 |
| | 0.003 | 832 (45.1%) | 72 (97.3%) | 48 (21.1%) | 581.8 | 566.1 | 1263.2 | 265.9 | 0.320 |
| | 0.0035 | 771 (41.8%) | 71 (95.9%) | 40 (17.5%) | 570.5 | 547.4 | 1274.5 | 223.6 | 0.290 |
| | 0.004 | 725 (39.3%) | 68 (91.9%) | 36 (15.8%) | 569.9 | 523.7 | 1275.1 | 201.3 | 0.278 |
| | 0.0045 | 680 (36.9%) | 68 (91.9%) | 31 (13.6%) | 548.1 | 503.7 | 1296.9 | 176.3 | 0.259 |
| | 0.005 | 653 (35.4%) | 67 (90.5%) | 27 (11.8%) | 552.1 | 499.9 | 1292.9 | 153.1 | 0.234 |
| | 0.0055 | 623 (33.8%) | 65 (87.8%) | 21 (9.2%) | 576.2 | 506.1 | 1268.8 | 116.9 | 0.188 |
| | 0.006 | 604 (32.7%) | 65 (87.8%) | 17 (7.5%) | 580.3 | 509.7 | 1264.7 | 94.3 | 0.156 |
| | 0.0065 | 583 (31.6%) | 65 (87.8%) | 15 (6.6%) | 568.1 | 499.0 | 1276.9 | 84.0 | 0.144 |
| | 0.007 | 563 (30.5%) | 65 (87.8%) | 13 (5.7%) | 557.4 | 489.6 | 1287.6 | 73.4 | 0.130 |
| | 0.008 | 538 (29.2%) | 65 (87.8%) | 11 (4.8%) | 540.9 | 475.1 | 1304.1 | 62.9 | 0.117 |
| | 0.009 | 514 (27.9%) | 62 (83.8%) | 10 (4.4%) | 545.5 | 457.0 | 1299.5 | 57.0 | 0.111 |
| **Relative ASV abundance by library** (Threshold = minimum proportion of read-abundance relative to the total number of reads in that library) | 0.01 | 498 (27%) | 58 (78.4%) | 8 (3.5%) | 578.7 | 453.6 | 1266.3 | 44.4 | 0.089 |
| | 0.015 | 431 (23.4%) | 53 (71.6%) | 3 (1.3%) | 578.5 | 414.3 | 1266.5 | 16.7 | 0.039 |
| | 0.02 | 387 (21%) | 48 (64.9%) | 2 (0.9%) | 579.5 | 375.9 | 1265.5 | 11.1 | 0.029 |
| | 0.025 | 352 (19.1%) | 44 (59.5%) | 1 (0.4%) | 582.7 | 346.5 | 1262.3 | 5.5 | 0.016 |
| | 0.03 | 326 (17.7%) | 42 (56.8%) | 1 (0.4%) | 564.5 | 320.4 | 1280.5 | 5.6 | 0.017 |
| | 0.035 | 302 (16.4%) | 39 (52.7%) | 1 (0.4%) | 562.4 | 296.4 | 1282.6 | 5.6 | 0.019 |
| | 0.04 | 289 (15.7%) | 38 (51.4%) | 1 (0.4%) | 551.7 | 283.3 | 1293.3 | 5.7 | 0.020 |
| | 0.045 | 265 (14.4%) | 36 (48.6%) | 0 (0%) | 544.7 | 265.0 | 1300.3 | 0.0 | 0.000 |
| | 0.05 | 245 (13.3%) | 32 (43.2%) | 0 (0%) | 566.6 | 245.0 | 1278.4 | 0.0 | 0.000 |
| | 0.055 | 236 (12.8%) | 31 (41.9%) | 0 (0%) | 563.4 | 236.0 | 1281.6 | 0.0 | 0.000 |
| | 0.06 | 222 (12%) | 31 (41.9%) | 0 (0%) | 529.9 | 222.0 | 1315.1 | 0.0 | 0.000 |
| | 0.065 | 212 (11.5%) | 29 (39.2%) | 0 (0%) | 541.0 | 212.0 | 1304.0 | 0.0 | 0.000 |
| | 0.075 | 193 (10.5%) | 27 (36.5%) | 0 (0%) | 529.0 | 193.0 | 1316.0 | 0.0 | 0.000 |
| | 0.09 | 177 (9.6%) | 26 (35.1%) | 0 (0%) | 503.8 | 177.0 | 1341.2 | 0.0 | 0.000 |
| | 0.1 | 168 (9.1%) | 26 (35.1%) | 0 (0%) | 478.2 | 168.0 | 1366.8 | 0.0 | 0.000 |
| | 0.15 | 119 (6.4%) | 19 (25.7%) | 0 (0%) | 463.5 | 119.0 | 1381.5 | 0.0 | 0.000 |
| | 0.2 | 100 (5.4%) | 16 (21.6%) | 0 (0%) | 462.5 | 100.0 | 1382.5 | 0.0 | 0.000 |
| | 0.25 | 81 (4.4%) | 14 (18.9%) | 0 (0%) | 428.1 | 81.0 | 1416.9 | 0.0 | 0.000 |
| | 0.3 | 66 (3.6%) | 14 (18.9%) | 0 (0%) | 348.9 | 66.0 | 1496.1 | 0.0 | 0.000 |
| | 0.35 | 55 (3%) | 13 (17.6%) | 0 (0%) | 313.1 | 55.0 | 1531.9 | 0.0 | 0.000 |
| | 0.4 | 48 (2.6%) | 13 (17.6%) | 0 (0%) | 273.2 | 48.0 | 1571.8 | 0.0 | 0.000 |

* Observed values; ** Estimated values; *** The ratio n/t represents the estimated proportion of na-ASVs in the filtered dataset

**Table S8.** Filtering performance of filtering criteria "Relative ASV abundance by library and 15% clade" for a range of minimum threshold values for read abundance for the COL dataset.

| Criterium | Thershold | Surviving ASVs (t)* | Surviving va-ASVs (av)* | Surviving vna-ASVs (nv)* | Initial a-ASVs (A)** | Surviving a-ASV (a)** | Initial na-ASVs (N)** | Surviving na-ASV (n)** | n/t *** |
|---|---|---|---|---|---|---|---|---|---|
| **Pre-filtering initial values** | | T = 1845 | Av = 74 | Nv = 228 | | | | | |
| **Relative ASV abundance by library and 15% clade** (Threshold = minimum read proportion relative to the total number of reads in the 15% divergence clade where each ASV is included in that library) | 0.001 | 1759 (95.3%) | 74 (100%) | 216 (94.7%) | 211.0 | 211.0 | 1634.0 | 1548.0 | 0.880 |
| | 0.0025 | 1598 (86.6%) | 74 (100%) | 185 (81.1%) | 535.3 | 535.3 | 1309.7 | 1062.7 | 0.665 |
| | 0.005 | 1406 (76.2%) | 74 (100%) | 142 (62.3%) | 681.1 | 681.1 | 1163.9 | 724.9 | 0.516 |
| | 0.01 | 1225 (66.4%) | 73 (98.6%) | 107 (46.9%) | 694.4 | 685.0 | 1150.6 | 540.0 | 0.441 |
| | 0.015 | 1122 (60.8%) | 73 (98.6%) | 90 (39.5%) | 665.3 | 656.3 | 1179.7 | 465.7 | 0.415 |
| | 0.02 | 1062 (57.6%) | 72 (97.3%) | 79 (34.6%) | 674.8 | 656.5 | 1170.2 | 405.5 | 0.382 |
| | 0.025 | 1014 (55%) | 72 (97.3%) | 75 (32.9%) | 632.1 | 615.0 | 1212.9 | 399.0 | 0.393 |
| | 0.03 | 965 (52.3%) | 72 (97.3%) | 70 (30.7%) | 598.5 | 582.3 | 1246.5 | 382.7 | 0.397 |
| | 0.035 | 939 (50.9%) | 72 (97.3%) | 67 (29.4%) | 584.3 | 568.5 | 1260.7 | 370.5 | 0.395 |
| | 0.04 | 924 (50.1%) | 72 (97.3%) | 66 (28.9%) | 570.5 | 555.1 | 1274.5 | 368.9 | 0.399 |
| | 0.045 | 899 (48.7%) | 72 (97.3%) | 61 (26.8%) | 574.7 | 559.1 | 1270.3 | 339.9 | 0.378 |
| | 0.05 | 866 (46.9%) | 72 (97.3%) | 57 (25%) | 559.8 | 544.7 | 1285.2 | 321.3 | 0.371 |
| | 0.055 | 852 (46.2%) | 72 (97.3%) | 57 (25%) | 540.5 | 525.9 | 1304.5 | 326.1 | 0.383 |
| | 0.06 | 839 (45.5%) | 72 (97.3%) | 56 (24.6%) | 530.5 | 516.1 | 1314.5 | 322.9 | 0.385 |
| | 0.065 | 824 (44.7%) | 72 (97.3%) | 53 (23.2%) | 533.6 | 519.2 | 1311.4 | 304.8 | 0.370 |
| | 0.075 | 802 (43.5%) | 72 (97.3%) | 52 (22.8%) | 511.8 | 497.9 | 1333.2 | 304.1 | 0.379 |
| | 0.09 | 764 (41.4%) | 71 (95.9%) | 46 (20.2%) | 517.0 | 496.1 | 1328.0 | 267.9 | 0.351 |
| | 0.1 | 751 (40.7%) | 71 (95.9%) | 44 (19.3%) | 515.3 | 494.4 | 1329.7 | 256.6 | 0.342 |
| | 0.15 | 669 (36.3%) | 70 (94.6%) | 35 (15.4%) | 486.8 | 460.5 | 1358.2 | 208.5 | 0.312 |
| | 0.2 | 624 (33.8%) | 70 (94.6%) | 32 (14%) | 453.1 | 428.7 | 1391.9 | 195.3 | 0.313 |
| | 0.25 | 581 (31.5%) | 70 (94.6%) | 31 (13.6%) | 407.6 | 385.6 | 1437.4 | 195.4 | 0.336 |
| | 0.3 | 557 (30.2%) | 70 (94.6%) | 29 (12.7%) | 393.7 | 372.4 | 1451.3 | 184.6 | 0.331 |
| | 0.35 | 519 (28.1%) | 68 (91.9%) | 25 (11%) | 391.3 | 359.6 | 1453.7 | 159.4 | 0.307 |
| | 0.4 | 505 (27.4%) | 68 (91.9%) | 25 (11%) | 374.0 | 343.7 | 1471.0 | 161.3 | 0.319 |
| | 0.5 | 470 (25.5%) | 67 (90.5%) | 23 (10.1%) | 352.9 | 319.5 | 1492.1 | 150.5 | 0.320 |
| | 0.6 | 426 (23.1%) | 65 (87.8%) | 18 (7.9%) | 350.7 | 308.0 | 1494.3 | 118.0 | 0.277 |
| | 0.7 | 401 (21.7%) | 62 (83.8%) | 17 (7.5%) | 345.1 | 289.2 | 1499.9 | 111.8 | 0.279 |
| | 0.8 | 378 (20.5%) | 62 (83.8%) | 16 (7%) | 323.7 | 271.2 | 1521.3 | 106.8 | 0.282 |
| | 0.9 | 329 (17.8%) | 53 (71.6%) | 15 (6.6%) | 319.2 | 228.6 | 1525.8 | 100.4 | 0.305 |

* Observed values; ** Estimated values; *** The ratio n/t represents the estimated proportion of na-ASVs in the filtered dataset

**Table S9.** Filtering performance of filtering criteria "Relative ASV abundance by library and 20% clade" for a range of minimum threshold values for read abundance for the COL dataset.

| Criterium | Thershold | Surviving ASVs (t)* | Surviving va-ASVs (av)* | Surviving vna-ASVs (nv)* | Initial a-ASVs (A)** | Surviving a-ASV (a)** | Initial na-ASVs (N)** | Surviving na-ASV (n)** | n/t *** |
|---|---|---|---|---|---|---|---|---|---|
| **Pre-filtering initial values** | | T = 1845 | Av = 74 | Nv = 228 | | | | | |
| **Relative ASV abundance by library and 20% clade** (Threshold = minimum read proportion relative to the total number of reads in the 20% divergence clade where each ASV is included in that library) | 0.001 | 1749 (94.8%) | 74 (100%) | 213 (93.4%) | 385.8 | 385.8 | 1459.2 | 1363.2 | 0.779 |
| | 0.0025 | 1509 (81.8%) | 74 (100%) | 158 (69.3%) | 750.6 | 750.6 | 1094.4 | 758.4 | 0.503 |
| | 0.005 | 1289 (69.9%) | 74 (100%) | 109 (47.8%) | 779.7 | 779.7 | 1065.3 | 509.3 | 0.395 |
| | 0.01 | 1077 (58.4%) | 72 (97.3%) | 70 (30.7%) | 766.6 | 745.9 | 1078.4 | 331.1 | 0.307 |
| | 0.015 | 960 (52%) | 71 (95.9%) | 50 (21.9%) | 750.4 | 719.9 | 1094.6 | 240.1 | 0.250 |
| | 0.02 | 897 (48.6%) | 70 (94.6%) | 42 (18.4%) | 731.4 | 691.9 | 1113.6 | 205.1 | 0.229 |
| | 0.025 | 852 (46.2%) | 69 (93.2%) | 38 (16.7%) | 711.1 | 663.0 | 1133.9 | 189.0 | 0.222 |
| | 0.03 | 808 (43.8%) | 69 (93.2%) | 32 (14%) | 693.2 | 646.3 | 1151.8 | 161.7 | 0.200 |
| | 0.035 | 772 (41.8%) | 69 (93.2%) | 27 (11.8%) | 680.0 | 634.0 | 1165.0 | 138.0 | 0.179 |
| | 0.04 | 750 (40.7%) | 68 (91.9%) | 27 (11.8%) | 664.0 | 610.1 | 1181.0 | 139.9 | 0.186 |
| | 0.045 | 729 (39.5%) | 66 (89.2%) | 22 (9.6%) | 692.7 | 617.8 | 1152.3 | 111.2 | 0.153 |
| | 0.05 | 708 (38.4%) | 66 (89.2%) | 20 (8.8%) | 679.2 | 605.7 | 1165.8 | 102.3 | 0.144 |
| | 0.055 | 695 (37.7%) | 64 (86.5%) | 20 (8.8%) | 686.0 | 593.3 | 1159.0 | 101.7 | 0.146 |
| | 0.06 | 680 (36.9%) | 64 (86.5%) | 19 (8.3%) | 673.4 | 582.4 | 1171.6 | 97.6 | 0.144 |
| | 0.065 | 663 (35.9%) | 64 (86.5%) | 16 (7%) | 671.4 | 580.6 | 1173.6 | 82.4 | 0.124 |
| | 0.075 | 644 (34.9%) | 63 (85.1%) | 16 (7%) | 658.7 | 560.7 | 1186.3 | 83.3 | 0.129 |
| | 0.09 | 613 (33.2%) | 62 (83.8%) | 14 (6.1%) | 643.6 | 539.2 | 1201.4 | 73.8 | 0.120 |
| | 0.1 | 600 (32.5%) | 62 (83.8%) | 13 (5.7%) | 633.7 | 530.9 | 1211.3 | 69.1 | 0.115 |
| | 0.15 | 529 (28.7%) | 59 (79.7%) | 11 (4.8%) | 587.4 | 468.3 | 1257.6 | 60.7 | 0.115 |
| | 0.2 | 492 (26.7%) | 58 (78.4%) | 10 (4.4%) | 555.6 | 435.4 | 1289.4 | 56.6 | 0.115 |
| | 0.25 | 452 (24.5%) | 56 (75.7%) | 10 (4.4%) | 520.5 | 393.9 | 1324.5 | 58.1 | 0.129 |
| | 0.3 | 425 (23%) | 56 (75.7%) | 8 (3.5%) | 499.2 | 377.8 | 1345.8 | 47.2 | 0.111 |
| | 0.35 | 390 (21.1%) | 53 (71.6%) | 7 (3.1%) | 486.3 | 348.3 | 1358.7 | 41.7 | 0.107 |
| | 0.4 | 379 (20.5%) | 53 (71.6%) | 7 (3.1%) | 470.2 | 336.8 | 1374.8 | 42.2 | 0.111 |
| | 0.5 | 355 (19.2%) | 52 (70.3%) | 7 (3.1%) | 444.0 | 312.0 | 1401.0 | 43.0 | 0.121 |
| | 0.6 | 319 (17.3%) | 48 (64.9%) | 5 (2.2%) | 444.4 | 288.3 | 1400.6 | 30.7 | 0.096 |
| | 0.7 | 296 (16%) | 46 (62.2%) | 5 (2.2%) | 426.1 | 264.9 | 1418.9 | 31.1 | 0.105 |
| | 0.8 | 275 (14.9%) | 46 (62.2%) | 5 (2.2%) | 391.1 | 243.1 | 1453.9 | 31.9 | 0.116 |
| | 0.9 | 238 (12.9%) | 38 (51.4%) | 5 (2.2%) | 401.8 | 206.4 | 1443.2 | 31.6 | 0.133 |

* Observed values; ** Estimated values; *** The ratio n/t represents the estimated proportion of na-ASVs in the filtered dataset

**Table S10.** Filtering performance of filtering criteria "Relative ASV abundance by library and 26% clade" for a range of minimum threshold values for read abundance for the COL dataset.

| Criterium | Thershold | Surviving ASVs (t)* | Surviving va-ASVs (av)* | Surviving vna-ASVs (nv)* | Initial a-ASVs (A)** | Surviving a-ASV (a)** | Initial na-ASVs (N)** | Surviving na-ASV (n)** | n/t *** |
|---|---|---|---|---|---|---|---|---|---|
| **Pre-filtering initial values** | | T = 1845 | Av = 74 | Nv = 228 | | | | | |
| **Relative ASV abundance by library and 26% clade** (Threshold = minimum read proportion relative to the total number of reads in the 26% divergence clade where each ASV is included in that library) | 0.001 | 1696 (91.9%) | 74 (100%) | 200 (87.7%) | 631.7 | 631.7 | 1213.3 | 1064.3 | 0.628 |
| | 0.0025 | 1374 (74.5%) | 74 (100%) | 133 (58.3%) | 714.6 | 714.6 | 1130.4 | 659.4 | 0.480 |
| | 0.005 | 1070 (58%) | 73 (98.6%) | 77 (33.8%) | 688.9 | 679.5 | 1156.1 | 390.5 | 0.365 |
| | 0.01 | 853 (46.2%) | 70 (94.6%) | 48 (21.1%) | 631.7 | 597.6 | 1213.3 | 255.4 | 0.299 |
| | 0.015 | 726 (39.3%) | 67 (90.5%) | 29 (12.7%) | 631.4 | 571.6 | 1213.6 | 154.4 | 0.213 |
| | 0.02 | 684 (37.1%) | 65 (87.8%) | 24 (10.5%) | 633.5 | 556.5 | 1211.5 | 127.5 | 0.186 |
| | 0.025 | 643 (34.9%) | 62 (83.8%) | 21 (9.2%) | 634.4 | 531.5 | 1210.6 | 111.5 | 0.173 |
| | 0.03 | 601 (32.6%) | 60 (81.1%) | 17 (7.5%) | 629.5 | 510.4 | 1215.5 | 90.6 | 0.151 |
| | 0.035 | 575 (31.2%) | 59 (79.7%) | 13 (5.7%) | 634.6 | 506.0 | 1210.4 | 69.0 | 0.120 |
| | 0.04 | 549 (29.8%) | 59 (79.7%) | 12 (5.3%) | 606.8 | 483.8 | 1238.2 | 65.2 | 0.119 |
| | 0.045 | 537 (29.1%) | 57 (77%) | 12 (5.3%) | 613.0 | 472.2 | 1232.0 | 64.8 | 0.121 |
| | 0.05 | 524 (28.4%) | 54 (73%) | 11 (4.8%) | 638.3 | 465.8 | 1206.7 | 58.2 | 0.111 |
| | 0.055 | 508 (27.5%) | 52 (70.3%) | 11 (4.8%) | 640.2 | 449.9 | 1204.8 | 58.1 | 0.114 |
| | 0.06 | 487 (26.4%) | 50 (67.6%) | 11 (4.8%) | 634.3 | 428.6 | 1210.7 | 58.4 | 0.120 |
| | 0.065 | 475 (25.7%) | 49 (66.2%) | 9 (3.9%) | 645.9 | 427.7 | 1199.1 | 47.3 | 0.100 |
| | 0.075 | 453 (24.6%) | 47 (63.5%) | 9 (3.9%) | 638.2 | 405.4 | 1206.8 | 47.6 | 0.105 |
| | 0.09 | 422 (22.9%) | 44 (59.5%) | 8 (3.5%) | 638.5 | 379.7 | 1206.5 | 42.3 | 0.100 |
| | 0.1 | 409 (22.2%) | 43 (58.1%) | 8 (3.5%) | 630.5 | 366.4 | 1214.5 | 42.6 | 0.104 |
| | 0.15 | 339 (18.4%) | 38 (51.4%) | 6 (2.6%) | 596.2 | 306.1 | 1248.8 | 32.9 | 0.097 |
| | 0.2 | 310 (16.8%) | 36 (48.6%) | 6 (2.6%) | 568.2 | 276.4 | 1276.8 | 33.6 | 0.108 |
| | 0.25 | 276 (15%) | 29 (39.2%) | 6 (2.6%) | 622.2 | 243.8 | 1222.8 | 32.2 | 0.117 |
| | 0.3 | 252 (13.7%) | 28 (37.8%) | 6 (2.6%) | 577.9 | 218.7 | 1267.1 | 33.3 | 0.132 |
| | 0.35 | 232 (12.6%) | 26 (35.1%) | 6 (2.6%) | 564.4 | 198.3 | 1280.6 | 33.7 | 0.145 |
| | 0.4 | 215 (11.7%) | 25 (33.8%) | 6 (2.6%) | 534.3 | 180.5 | 1310.7 | 34.5 | 0.160 |
| | 0.5 | 191 (10.4%) | 23 (31.1%) | 6 (2.6%) | 500.7 | 155.6 | 1344.3 | 35.4 | 0.185 |
| | 0.6 | 162 (8.8%) | 20 (27%) | 4 (1.8%) | 512.9 | 138.6 | 1332.1 | 23.4 | 0.144 |
| | 0.7 | 144 (7.8%) | 18 (24.3%) | 3 (1.3%) | 520.3 | 126.6 | 1324.7 | 17.4 | 0.121 |
| | 0.8 | 128 (6.9%) | 16 (21.6%) | 3 (1.3%) | 510.8 | 110.4 | 1334.2 | 17.6 | 0.137 |
| | 0.9 | 104 (5.6%) | 14 (18.9%) | 3 (1.3%) | 452.9 | 85.7 | 1392.1 | 18.3 | 0.176 |

* Observed values; ** Estimated values; *** The ratio n/t represents the estimated proportion of na-ASVs in the filtered dataset

#### 2.3. Supplementary tables for the CAC Dataset (Tables S11-S15)

**Table S11.** Filtering performance of filtering criteria "Absolute ASV abundance by library" for a range of minimum threshold values for read abundance for the CAC dataset.

| Criterium | Thershold | Surviving ASVs (t)* | Surviving va-ASVs (av)* | Surviving vna-ASVs (nv)* | Initial a-ASVs (A)** | Surviving a-ASV (a)** | Initial na-ASVs (N)** | Surviving na-ASV (n)** | n/t *** |
|---|---|---|---|---|---|---|---|---|---|
| **Pre-filtering initial values** | | **T = 4804** | **Av = 221** | **Nv = 237** | | | | | |
| **Absolute ASV abundance by library** (Threshold = minimum threshold of read-abundance in a given library) | 3 | 3165 (65.9%) | 219 (99.1%) | 89 (37.6%) | 2211.4 | 2191.4 | 2592.6 | 973.6 | 0.31 |
| | 4 | 2758 (57.4%) | 213 (96.4%) | 61 (25.7%) | 2153.9 | 2075.9 | 2650.1 | 682.1 | 0.25 |
| | 5 | 2531 (52.7%) | 212 (95.9%) | 52 (21.9%) | 1996.2 | 1915.0 | 2807.8 | 616.0 | 0.24 |
| | 8 | 2126 (44.3%) | 202 (91.4%) | 34 (14.3%) | 1864.6 | 1704.3 | 2939.4 | 421.7 | 0.20 |
| | 10 | 1953 (40.7%) | 196 (88.7%) | 22 (9.3%) | 1897.9 | 1683.2 | 2906.1 | 269.8 | 0.14 |
| | 12 | 1801 (37.5%) | 192 (86.9%) | 17 (7.2%) | 1827.3 | 1587.5 | 2976.7 | 213.5 | 0.12 |
| | 15 | 1635 (34%) | 186 (84.2%) | 15 (6.3%) | 1710.0 | 1439.2 | 3094.0 | 195.8 | 0.12 |
| | 20 | 1432 (29.8%) | 178 (80.5%) | 10 (4.2%) | 1610.6 | 1297.3 | 3193.4 | 134.7 | 0.09 |
| | 25 | 1299 (27%) | 167 (75.6%) | 9 (3.8%) | 1555.8 | 1175.7 | 3248.2 | 123.3 | 0.09 |
| | 35 | 1086 (22.6%) | 157 (71%) | 7 (3%) | 1386.6 | 985.1 | 3417.4 | 100.9 | 0.09 |
| | 40 | 1022 (21.3%) | 155 (70.1%) | 5 (2.1%) | 1353.4 | 949.2 | 3450.6 | 72.8 | 0.07 |
| | 45 | 967 (20.1%) | 147 (66.5%) | 5 (2.1%) | 1344.0 | 894.0 | 3460.0 | 73.0 | 0.08 |
| | 50 | 918 (19.1%) | 145 (65.6%) | 5 (2.1%) | 1286.0 | 843.8 | 3518.0 | 74.2 | 0.08 |
| | 60 | 816 (17%) | 138 (62.4%) | 5 (2.1%) | 1184.5 | 739.6 | 3619.5 | 76.4 | 0.09 |
| | 70 | 740 (15.4%) | 135 (61.1%) | 4 (1.7%) | 1109.3 | 677.6 | 3694.7 | 62.4 | 0.08 |
| | 80 | 678 (14.1%) | 126 (57%) | 4 (1.7%) | 1078.9 | 615.1 | 3725.1 | 62.9 | 0.09 |
| | 90 | 630 (13.1%) | 120 (54.3%) | 4 (1.7%) | 1043.4 | 566.5 | 3760.6 | 63.5 | 0.10 |
| | 100 | 588 (12.2%) | 117 (52.9%) | 2 (0.8%) | 1050.8 | 556.3 | 3753.2 | 31.7 | 0.05 |

* Observed values; ** Estimated values; *** The ratio n/t represents the estimated proportion of na-ASVs in the filtered dataset.

**Table S12.** Filtering performance of filtering criteria "Relative ASV abundance by library" for a range of minimum threshold values for read abundance for the CAC dataset.

| Criterium | Thershold | Surviving ASVs (t)* | Surviving va-ASVs (av)* | Surviving vna-ASVs (nv)* | Initial a-ASVs (A)** | Surviving a-ASV (a)** | Initial na-ASVs (N)** | Surviving na-ASV (n)** | n/t *** |
|---|---|---|---|---|---|---|---|---|---|
| **Pre-filtering initial values** | | T = 4804 | Av = 221 | Nv = 237 | | | | | |
| **Relative ASV abundance by library** (Threshold = minimum proportion of read-abundance relative to the total number of reads in that library) | 0.00025 | 3344 (69.6%) | 217 (98.2%) | 98 (41.4%) | 2388.4 | 2345.1 | 2415.6 | 998.9 | 0.30 |
| | 0.0005 | 2521 (52.5%) | 211 (95.5%) | 53 (22.4%) | 1978.7 | 1889.2 | 2825.3 | 631.8 | 0.25 |
| | 0.001 | 1944 (40.5%) | 199 (90%) | 26 (11%) | 1791.9 | 1613.6 | 3012.1 | 330.4 | 0.17 |
| | 0.0015 | 1646 (34.3%) | 188 (85.1%) | 17 (7.2%) | 1670.7 | 1421.3 | 3133.3 | 224.7 | 0.14 |
| | 0.002 | 1445 (30.1%) | 180 (81.4%) | 12 (5.1%) | 1573.3 | 1281.4 | 3230.7 | 163.6 | 0.11 |
| | 0.0025 | 1309 (27.2%) | 175 (79.2%) | 8 (3.4%) | 1512.8 | 1197.9 | 3291.2 | 111.1 | 0.08 |
| | 0.003 | 1187 (24.7%) | 164 (74.2%) | 6 (2.5%) | 1486.4 | 1103.0 | 3317.6 | 84.0 | 0.07 |
| | 0.0035 | 1092 (22.7%) | 157 (71%) | 6 (2.5%) | 1416.4 | 1006.2 | 3387.6 | 85.8 | 0.08 |
| | 0.004 | 1007 (21%) | 153 (69.2%) | 4 (1.7%) | 1370.9 | 949.1 | 3433.1 | 57.9 | 0.06 |
| | 0.0045 | 943 (19.6%) | 148 (67%) | 4 (1.7%) | 1320.3 | 884.2 | 3483.7 | 58.8 | 0.06 |
| | 0.005 | 904 (18.8%) | 147 (66.5%) | 4 (1.7%) | 1269.4 | 844.3 | 3534.6 | 59.7 | 0.07 |
| | 0.0055 | 867 (18%) | 146 (66.1%) | 4 (1.7%) | 1220.8 | 806.5 | 3583.2 | 60.5 | 0.07 |
| | 0.006 | 827 (17.2%) | 141 (63.8%) | 4 (1.7%) | 1200.9 | 766.2 | 3603.1 | 60.8 | 0.07 |
| | 0.0065 | 781 (16.3%) | 135 (61.1%) | 4 (1.7%) | 1178.4 | 719.8 | 3625.6 | 61.2 | 0.08 |
| | 0.007 | 746 (15.5%) | 132 (59.7%) | 3 (1.3%) | 1172.0 | 700.0 | 3632.0 | 46.0 | 0.06 |
| | 0.008 | 690 (14.4%) | 126 (57%) | 3 (1.3%) | 1128.6 | 643.5 | 3675.4 | 46.5 | 0.07 |
| | 0.009 | 643 (13.4%) | 124 (56.1%) | 3 (1.3%) | 1061.6 | 595.6 | 3742.4 | 47.4 | 0.07 |
| | 0.01 | 601 (12.5%) | 119 (53.8%) | 2 (0.8%) | 1057.4 | 569.4 | 3746.6 | 31.6 | 0.05 |

* Observed values; ** Estimated values; *** The ratio n/t represents the estimated proportion of na-ASVs in the filtered dataset

**Table S13.** Filtering performance of filtering criteria "Relative ASV abundance by library and 15% clade" for a range of minimum threshold values for read abundance for the CAC dataset.

| Criterium | Thershold | Surviving ASVs (t)* | Surviving va-ASVs (av)* | Surviving vna-ASVs (nv)* | Initial a-ASVs (A)** | Surviving a-ASV (a)** | Initial na-ASVs (N)** | Surviving na-ASV (n)** | n/t *** |
|---|---|---|---|---|---|---|---|---|---|
| **Pre-filtering initial values** | | T = 4804 | Av = 221 | Nv = 237 | | | | | |
| **Relative ASV abundance by library and 15% clade** (Threshold = minimum read proportion relative to the total number of reads in the 15% divergence clade where each ASV is included in that library) | 0.001 | 4659 (97%) | 221 (100%) | 217 (91.6%) | 3085.8 | 3085.8 | 1718.3 | 1573.3 | 0.34 |
| | 0.0025 | 4446 (92.5%) | 221 (100%) | 193 (81.4%) | 2875.7 | 2875.7 | 1928.3 | 1570.3 | 0.35 |
| | 0.005 | 4187 (87.2%) | 221 (100%) | 158 (66.7%) | 2953.0 | 2953.0 | 1851.0 | 1234.0 | 0.29 |
| | 0.01 | 3908 (81.3%) | 220 (99.5%) | 128 (54%) | 2884.2 | 2871.1 | 1919.8 | 1036.9 | 0.27 |
| | 0.015 | 3674 (76.5%) | 220 (99.5%) | 106 (44.7%) | 2782.4 | 2769.8 | 2021.6 | 904.2 | 0.25 |
| | 0.02 | 3520 (73.3%) | 220 (99.5%) | 95 (40.1%) | 2681.2 | 2669.1 | 2122.8 | 850.9 | 0.24 |
| | 0.025 | 3402 (70.8%) | 220 (99.5%) | 86 (36.3%) | 2622.1 | 2610.3 | 2181.9 | 791.7 | 0.23 |
| | 0.03 | 3298 (68.7%) | 219 (99.1%) | 77 (32.5%) | 2608.2 | 2584.6 | 2195.8 | 713.4 | 0.22 |
| | 0.035 | 3196 (66.5%) | 219 (99.1%) | 70 (29.5%) | 2554.8 | 2531.7 | 2249.2 | 664.3 | 0.21 |
| | 0.04 | 3116 (64.9%) | 218 (98.6%) | 66 (27.8%) | 2511.7 | 2477.7 | 2292.3 | 638.3 | 0.20 |
| | 0.045 | 3048 (63.4%) | 217 (98.2%) | 63 (26.6%) | 2473.2 | 2428.4 | 2330.8 | 619.6 | 0.20 |
| | 0.05 | 2984 (62.1%) | 217 (98.2%) | 59 (24.9%) | 2439.5 | 2395.4 | 2364.5 | 588.6 | 0.20 |
| | 0.055 | 2921 (60.8%) | 217 (98.2%) | 57 (24.1%) | 2381.5 | 2338.4 | 2422.5 | 582.6 | 0.20 |
| | 0.06 | 2870 (59.7%) | 216 (97.7%) | 55 (23.2%) | 2354.9 | 2301.7 | 2449.1 | 568.3 | 0.20 |
| | 0.065 | 2827 (58.8%) | 216 (97.7%) | 55 (23.2%) | 2297.2 | 2245.3 | 2506.8 | 581.7 | 0.21 |
| | 0.075 | 2745 (57.1%) | 216 (97.7%) | 54 (22.8%) | 2201.9 | 2152.1 | 2602.1 | 592.9 | 0.22 |
| | 0.09 | 2627 (54.7%) | 216 (97.7%) | 45 (19%) | 2177.6 | 2128.3 | 2626.4 | 498.7 | 0.19 |
| | 0.1 | 2569 (53.5%) | 216 (97.7%) | 42 (17.7%) | 2146.6 | 2098.1 | 2657.4 | 470.9 | 0.18 |
| | 0.15 | 2250 (46.8%) | 209 (94.6%) | 33 (13.9%) | 1960.5 | 1854.1 | 2843.5 | 395.9 | 0.18 |
| | 0.2 | 2069 (43.1%) | 204 (92.3%) | 27 (11.4%) | 1880.6 | 1736.0 | 2923.4 | 333.0 | 0.16 |
| | 0.25 | 1923 (40%) | 202 (91.4%) | 25 (10.5%) | 1751.6 | 1601.0 | 3052.4 | 322.0 | 0.17 |
| | 0.3 | 1789 (37.2%) | 194 (87.8%) | 25 (10.5%) | 1660.2 | 1457.4 | 3143.8 | 331.6 | 0.19 |
| | 0.35 | 1692 (35.2%) | 190 (86%) | 21 (8.9%) | 1642.2 | 1411.8 | 3161.8 | 280.2 | 0.17 |
| | 0.4 | 1642 (34.2%) | 185 (83.7%) | 21 (8.9%) | 1625.0 | 1360.3 | 3179.0 | 281.7 | 0.17 |
| | 0.5 | 1514 (31.5%) | 176 (79.6%) | 18 (7.6%) | 1595.0 | 1270.2 | 3210.0 | 243.8 | 0.16 |
| | 0.6 | 1392 (29%) | 173 (78.3%) | 16 (6.8%) | 1492.5 | 1168.3 | 3313.5 | 223.7 | 0.16 |
| | 0.7 | 1291 (26.9%) | 162 (73.3%) | 10 (4.2%) | 1575.2 | 1154.6 | 3231.8 | 136.4 | 0.11 |
| | 0.8 | 1208 (25.1%) | 153 (69.2%) | 10 (4.2%) | 1546.1 | 1070.4 | 3261.9 | 137.6 | 0.11 |
| | 0.9 | 1083 (22.5%) | 142 (64.3%) | 8 (3.4%) | 1512.3 | 971.7 | 3296.7 | 111.3 | 0.10 |

* Observed values; ** Estimated values; *** The ratio n/t represents the estimated proportion of na-ASVs in the filtered dataset

**Table S14.** Filtering performance of filtering criteria "Relative ASV abundance by library and 20% clade" for a range of minimum threshold values for read abundance for the CAC dataset.

| Criterium | Thershold | Surviving ASVs (t)* | Surviving va-ASVs (av)* | Surviving vna-ASVs (nv)* | Initial a-ASVs (A)** | Surviving a-ASV (a)** | Initial na-ASVs (N)** | Surviving na-ASV (n)** | n/t *** |
|---|---|---|---|---|---|---|---|---|---|
| **Pre-filtering initial values** | | T = 4804 | Av = 221 | Nv = 237 | | | | | |
| **Relative ASV abundance by library and 20% clade** (Threshold = minimum read proportion relative to the total number of reads in the 20% divergence clade where each ASV is included in that library) | 0.001 | 4648 (96.8%) | 221 (100%) | 216 (91.1%) | 3043.4 | 3043.4 | 1760.6 | 1604.6 | 0.35 |
| | 0.0025 | 4412 (91.8%) | 221 (100%) | 189 (79.7%) | 2868.5 | 2868.5 | 1935.5 | 1543.5 | 0.35 |
| | 0.005 | 4116 (85.7%) | 221 (100%) | 148 (62.4%) | 2971.9 | 2971.9 | 1832.1 | 1144.1 | 0.28 |
| | 0.01 | 3786 (78.8%) | 220 (99.5%) | 119 (50.2%) | 2784.7 | 2772.1 | 2019.3 | 1013.9 | 0.27 |
| | 0.015 | 3547 (73.8%) | 220 (99.5%) | 97 (40.9%) | 2696.7 | 2684.5 | 2107.3 | 862.5 | 0.24 |
| | 0.02 | 3393 (70.6%) | 219 (99.1%) | 86 (36.3%) | 2626.7 | 2602.9 | 2177.3 | 790.1 | 0.23 |
| | 0.025 | 3265 (68%) | 219 (99.1%) | 76 (32.1%) | 2572.8 | 2549.5 | 2231.2 | 715.5 | 0.22 |
| | 0.03 | 3164 (65.9%) | 218 (98.6%) | 68 (28.7%) | 2552.7 | 2518.1 | 2251.3 | 645.9 | 0.20 |
| | 0.035 | 3049 (63.5%) | 218 (98.6%) | 60 (25.3%) | 2499.5 | 2465.6 | 2304.5 | 583.4 | 0.19 |
| | 0.04 | 2972 (61.9%) | 218 (98.6%) | 57 (24.1%) | 2435.4 | 2402.3 | 2368.6 | 569.7 | 0.19 |
| | 0.045 | 2904 (60.4%) | 217 (98.2%) | 53 (22.4%) | 2413.0 | 2369.3 | 2391.0 | 534.7 | 0.18 |
| | 0.05 | 2842 (59.2%) | 217 (98.2%) | 49 (20.7%) | 2385.0 | 2341.9 | 2419.0 | 500.1 | 0.18 |
| | 0.055 | 2784 (58%) | 217 (98.2%) | 47 (19.8%) | 2337.1 | 2294.8 | 2466.9 | 489.2 | 0.18 |
| | 0.06 | 2726 (56.7%) | 215 (97.3%) | 45 (19%) | 2316.6 | 2253.7 | 2487.4 | 472.3 | 0.17 |
| | 0.065 | 2686 (55.9%) | 215 (97.3%) | 44 (18.6%) | 2279.1 | 2217.2 | 2524.9 | 468.8 | 0.17 |
| | 0.075 | 2603 (54.2%) | 214 (96.8%) | 43 (18.1%) | 2200.3 | 2130.6 | 2603.7 | 472.4 | 0.18 |
| | 0.09 | 2480 (51.6%) | 213 (96.4%) | 35 (14.8%) | 2169.5 | 2090.9 | 2634.5 | 389.1 | 0.16 |
| | 0.1 | 2426 (50.5%) | 212 (95.9%) | 32 (13.5%) | 2156.3 | 2068.5 | 2647.7 | 357.5 | 0.15 |
| | 0.15 | 2095 (43.6%) | 203 (91.9%) | 23 (9.7%) | 1982.7 | 1821.2 | 2821.3 | 273.8 | 0.13 |
| | 0.2 | 1928 (40.1%) | 198 (89.6%) | 18 (7.6%) | 1906.3 | 1707.9 | 2897.7 | 220.1 | 0.11 |
| | 0.25 | 1788 (37.2%) | 195 (88.2%) | 17 (7.2%) | 1780.6 | 1571.1 | 3023.4 | 216.9 | 0.12 |
| | 0.3 | 1660 (34.6%) | 184 (83.3%) | 17 (7.2%) | 1728.9 | 1439.4 | 3075.1 | 220.6 | 0.13 |
| | 0.35 | 1559 (32.5%) | 180 (81.4%) | 14 (5.9%) | 1688.1 | 1374.9 | 3115.9 | 184.1 | 0.12 |
| | 0.4 | 1508 (31.4%) | 176 (79.6%) | 14 (5.9%) | 1660.3 | 1322.2 | 3144.7 | 185.8 | 0.12 |
| | 0.5 | 1376 (28.6%) | 166 (75.1%) | 10 (4.2%) | 1655.0 | 1243.1 | 3150.0 | 132.9 | 0.10 |
| | 0.6 | 1248 (26%) | 158 (71.5%) | 10 (4.2%) | 1553.7 | 1110.8 | 3252.3 | 137.2 | 0.11 |
| | 0.7 | 1152 (24%) | 149 (67.4%) | 4 (1.7%) | 1629.1 | 1098.4 | 3177.9 | 53.6 | 0.05 |
| | 0.8 | 1066 (22.2%) | 139 (62.9%) | 4 (1.7%) | 1609.0 | 1012.0 | 3199.0 | 54.0 | 0.05 |
| | 0.9 | 953 (19.8%) | 128 (57.9%) | 3 (1.3%) | 1574.7 | 912.1 | 3234.3 | 40.9 | 0.04 |

* Observed values; ** Estimated values; *** The ratio n/t represents the estimated proportion of na-ASVs in the filtered dataset

**Table S15.** Filtering performance of filtering criteria "Relative ASV abundance by library and 26% clade" for a range of minimum threshold values for read abundance for the CAC dataset.

| Criterium | Thershold | Surviving ASVs (t)* | Surviving va-ASVs (av)* | Surviving vna-ASVs (nv)* | Initial a-ASVs (A)** | Surviving a-ASV (a)** | Initial na-ASVs (N)** | Surviving na-ASV (n)** | n/t *** |
|---|---|---|---|---|---|---|---|---|---|
| **Pre-filtering initial values** | | T = 4804 | Av = 221 | Nv = 237 | | | | | |
| **Relative ASV abundance by library and 26% clade** (Threshold = minimum read proportion relative to the total number of reads in the 26% divergence clade where each ASV is included in that library) | 0.001 | 4315 (89.8%) | 221 (100%) | 191 (80.6%) | 2284.6 | 2284.6 | 2519.4 | 2030.4 | 0.47 |
| | 0.0025 | 3809 (79.3%) | 219 (99.1%) | 146 (61.6%) | 2266.0 | 2245.5 | 2538.0 | 1563.5 | 0.41 |
| | 0.005 | 3431 (71.4%) | 216 (97.7%) | 107 (45.1%) | 2399.9 | 2345.6 | 2404.1 | 1085.4 | 0.32 |
| | 0.01 | 2990 (62.2%) | 211 (95.5%) | 83 (35%) | 2162.9 | 2065.1 | 2641.1 | 924.9 | 0.31 |
| | 0.015 | 2720 (56.6%) | 208 (94.1%) | 64 (27%) | 2119.9 | 1995.2 | 2684.1 | 724.8 | 0.27 |
| | 0.02 | 2542 (52.9%) | 204 (92.3%) | 54 (22.8%) | 2081.9 | 1921.8 | 2722.1 | 620.2 | 0.24 |
| | 0.025 | 2383 (49.6%) | 201 (91%) | 46 (19.4%) | 2027.6 | 1844.1 | 2776.4 | 538.9 | 0.23 |
| | 0.03 | 2285 (47.6%) | 199 (90%) | 44 (18.6%) | 1949.0 | 1755.0 | 2855.0 | 530.0 | 0.23 |
| | 0.035 | 2174 (45.3%) | 198 (89.6%) | 37 (15.6%) | 1924.8 | 1724.5 | 2879.2 | 449.5 | 0.21 |
| | 0.04 | 2108 (43.9%) | 196 (88.7%) | 35 (14.8%) | 1892.0 | 1678.0 | 2912.0 | 430.0 | 0.20 |
| | 0.045 | 2051 (42.7%) | 195 (88.2%) | 34 (14.3%) | 1843.1 | 1626.2 | 2960.9 | 424.8 | 0.21 |
| | 0.05 | 1999 (41.6%) | 192 (86.9%) | 33 (13.9%) | 1823.2 | 1584.0 | 2980.8 | 415.0 | 0.21 |
| | 0.055 | 1937 (40.3%) | 191 (86.4%) | 32 (13.5%) | 1766.7 | 1526.9 | 3037.3 | 410.1 | 0.21 |
| | 0.06 | 1887 (39.3%) | 188 (85.1%) | 32 (13.5%) | 1730.4 | 1472.0 | 3073.6 | 415.0 | 0.22 |
| | 0.065 | 1844 (38.4%) | 186 (84.2%) | 31 (13.1%) | 1710.2 | 1439.3 | 3093.8 | 404.7 | 0.22 |
| | 0.075 | 1763 (36.7%) | 184 (83.3%) | 29 (12.2%) | 1654.7 | 1377.6 | 3149.3 | 385.4 | 0.22 |
| | 0.09 | 1668 (34.7%) | 183 (82.8%) | 23 (9.7%) | 1644.0 | 1361.3 | 3160.0 | 306.7 | 0.18 |
| | 0.1 | 1621 (33.7%) | 175 (79.2%) | 22 (9.3%) | 1681.0 | 1331.1 | 3123.0 | 289.9 | 0.18 |
| | 0.15 | 1400 (29.1%) | 160 (72.4%) | 18 (7.6%) | 1597.4 | 1156.5 | 3206.6 | 243.5 | 0.17 |
| | 0.2 | 1265 (26.3%) | 150 (67.9%) | 15 (6.3%) | 1561.4 | 1059.8 | 3242.6 | 205.2 | 0.16 |
| | 0.25 | 1158 (24.1%) | 142 (64.3%) | 14 (5.9%) | 1498.3 | 962.7 | 3305.7 | 195.3 | 0.17 |
| | 0.3 | 1054 (21.9%) | 136 (61.5%) | 14 (5.9%) | 1384.5 | 852.0 | 3419.5 | 202.0 | 0.19 |
| | 0.35 | 969 (20.2%) | 126 (57%) | 11 (4.6%) | 1424.5 | 812.1 | 3379.5 | 156.9 | 0.16 |
| | 0.4 | 924 (19.2%) | 117 (52.9%) | 11 (4.6%) | 1451.4 | 768.4 | 3352.6 | 155.6 | 0.17 |
| | 0.5 | 820 (17.1%) | 102 (46.2%) | 8 (3.4%) | 1537.7 | 709.7 | 3267.3 | 110.3 | 0.13 |
| | 0.6 | 738 (15.4%) | 93 (42.1%) | 8 (3.4%) | 1487.6 | 626.0 | 3318.4 | 112.0 | 0.15 |
| | 0.7 | 667 (13.9%) | 86 (38.9%) | 4 (1.7%) | 1573.8 | 612.4 | 3233.2 | 54.6 | 0.08 |
| | 0.8 | 607 (12.6%) | 76 (34.4%) | 4 (1.7%) | 1608.0 | 553.0 | 3200.0 | 54.0 | 0.09 |
| | 0.9 | 535 (11.1%) | 63 (28.5%) | 3 (1.3%) | 1740.5 | 496.2 | 3068.5 | 38.8 | 0.07 |

* Observed values; ** Estimated values; *** The ratio n/t represents the estimated proportion of na-ASVs in the filtered dataset

**Figure S1.** Schematic representation summarising the key steps in generating the amplicon sequence variant (ASV) datasets and NUMT dumping workflow. (**A)** Generation of BEE dataset; (**B)** Generation of COL and CAC datasets. (C) NUMT dumping workflow for the evaluation of alternative filtering criteria based on the survival/removal of verified authentic ASVs (va-ASVs; validated against a reference sequence) and verified non-authentic ASVs (vna-ASVs; those including STOP codons or INDELS). u-ASVs correspond to unclassified ASVs.

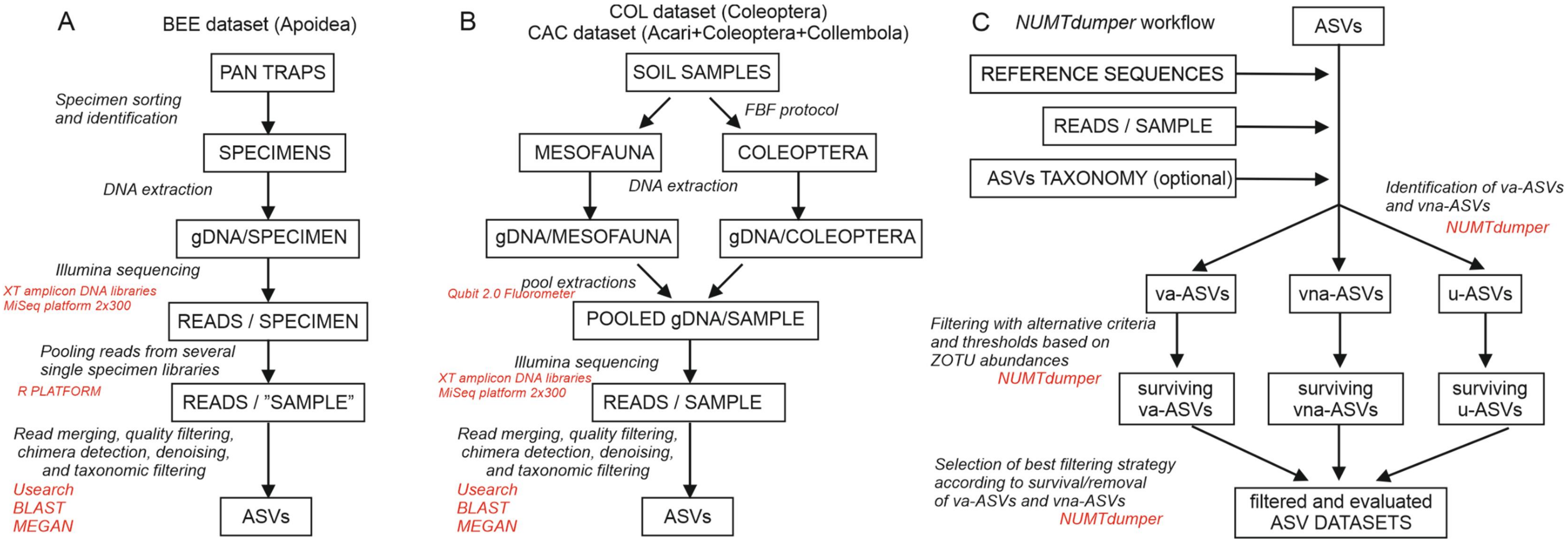
